# Shifting Perspectives on Biotherapeutic Treatment of Ulcerative Colitis using Lipid Mesophases: Formulation Design and Preclinical Validation

**DOI:** 10.64898/2026.02.09.701738

**Authors:** Gazzi Rafaela, Cremona Tiziana, Czekalla Reto, Campbell Emily, Schwarzfischer Marlene, Rogler Gerhard, Scharl Michael, Bergadano Alessandra, Mezzenga Raffaele, Kuentz Martin, Luciani Paola, Aleandri Simone

**Affiliations:** Department of Chemistry, Biochemistry and Pharmaceutical Sciences, University of Bern, Bern, Switzerland; Department of Gastroenterology and Hepatology, University Hospital Zurich, University of Zurich, Zurich, Switzerland; Experimental Animal Center (EAC), University of Bern, Bern, Switzerland; Laboratory of Food & Soft Materials, Institute of Food, Nutrition and Health, IFNH; Department for Health Sciences and Technology, D-HEST, ETH Zurich, Switzerland; Pharma Technology and Biotechnology. University of Applied Sciences and Arts Northwestern Switzerland, Muttenz, Switzerland

**Keywords:** Ulcerative colitis, colon-targeted drug delivery system, localized delivery of biotherapeutics, infliximab, lipid mesophases

## Abstract

Biotherapeutics are required to achieve high remission rates in patients with severe ulcerative colitis (UC); however, adverse effects, complex dosing regimens, administration routes, and low patient compliance may limit their widespread clinical use. Given the localized nature of UC, this study aimed to develop and evaluate a localized delivery strategy for infliximab (IFX), an anti-tumor necrosis factor-α (TNF-α) monoclonal antibody (mAb) recommended by the European Crohn’s and Colitis Organization (ECCO) and the American Crohn’s & Colitis Foundation for moderately-to-severely active UC. Exploiting the intrinsic biocompatibility, mucoadhesivity, and protein-entrapment capacity of lipid mesophases (LMPs), IFX was encapsulated within the gel matrix, providing protection against enzymatic and environmental degradation. IFX-loaded LMPs were designed for targeted delivery to inflamed colonic tissues via rectal or oral administration, with patient-centric oral dosage forms manufactured using a 3D printing approach. A comprehensive physicochemical characterization was performed to elucidate mesophase self-assembly and its relationship with IFX release profiles in biorelevant fluids. Therapeutic efficacy was evaluated in vivo using a dextran sulfate sodium (DSS)-induced colitis rat model, which demonstrated rectal gel retention for at least 8 h and colonic targeting of the oral formulation within 6 h. Under severe inflammatory conditions, LMP-based formulations reduced disease activity, inflammatory biomarkers (TNF-α and fecal lactoferrin), and colon shortening to values comparable to those of healthy controls, outperforming the therapeutic efficacy of subcutaneous IFX. Overall, this study establishes a biocompatible delivery platform that enables targeted colonic IFX release and suppresses systemic absorption, representing a promising advancement in the biotherapeutic treatment of UC.

## Introduction

Ulcerative colitis (UC) is a chronic inflammatory disorder of the colonic mucosa characterized by continuous inflammation and ulceration. Its pathogenesis involves a complex interplay between genetic and environmental factors (1–4), leading to symptoms that severely impair patients’ quality of life (1,2). UC typically begins in the rectum (proctitis) and may progress proximally to involve the entire colon (pancolitis) (1,2). Disease location and severity guide treatment selection (1,5), which includes salicylates, corticosteroids, calcineurin inhibitors, thiopurines, Janus kinase inhibitors and biotherapeutics such as mAbs (1,2).

Biologic therapies are recommended within a step-up approach for patients who fail to respond adequately to corticosteroids (6) and mainly include IFX (TNF-α), vedolizumab (anti-α4β7 integrin), and ustekinumab (anti-IL-12/IL-23), all of which demonstrate comparable clinical outcomes (7–11). Among these, IFX remains one of the most prescribed medications and is recommended by ECCO guidelines for moderately-to-severely active UC (6). IFX demonstrates superior efficacy in inducing clinical (12) and endoscopic remission (13) compared with other mAbs, and it is cost-effective during the first two years following diagnosis (14). Nevertheless, its clinical effectiveness may be compromised by immunogenicity, systemic toxicity, and treatment-related adverse events (15–17), with infusion-related reactions being the most frequently reported (5). Moreover, loss of response to biologics remains a major challenge, as up to 40% of patients in clinical remission develop anti-drug antibodies during IFX therapy (5,18). Consequently, the therapeutic benefits of IFX must be balanced against its adverse effects.

Because UC is confined to the colon, targeted drug delivery strategies are particularly attractive as they enable high mAb concentrations at sites of inflammation, thereby minimizing exposure to healthy tissue and reducing immunogenicity (19). Localized delivery can be achieved through rectal administration for distal UC (proctitis) or oral administration for extensive UC (pancolitis), potentially mitigating both anti-drug antibody formation and infusion-related reactions. However, colonic drug delivery poses route-dependent formulation challenges: conventional rectal drug formulations often exhibit poor mucoadhesion and leakage, while oral formulations must protect the drug during gastric transit and enable colonic release. To address these challenges, bioadhesive rectal (20) and gastro-resistant oral (21,22) protein-based formulations have been proposed.

Within this context, LMPs represent an attractive alternative drug delivery system, consisting of biodegradable gels capable of encapsulating hydrophilic, lipophilic and amphiphilic molecules. Proteins can be immobilized within the mesophase matrix, preventing enzymatic and environmental degradation while enabling controlled release (23). The safety and industrial translatability of LMPs are supported by the FluidCrystal^®^ technology developed by Camurus, a subcutaneous in situ-forming LMP gel. Moreover, we have previously demonstrated the efficacy of monolinolein (MLO)-based LMPs as a local treatment for UC (24). Upon hydration, MLO can form various structures, including highly viscous inverse cubic phases, whose appearance and rheology resemble cross-linked polymeric hydrogels (25). Although these structures can stabilize sensitive biomacromolecules, their narrow aqueous channels (approximately 3 nm) restrict the diffusion and release of large molecules such as mAbs (23,26,27). Previous studies have addressed this limitation by enlarging water channels through hydration-modulating agents (28,29) or charged lipids (27,30–32).

In this study, the hydration-modulating agent sucrose stearate (SE) was incorporated into MLO-based LMPs to engineer a localized mAb delivery system. Beyond its swelling properties (28), SE modulates LMP erosion in colonic fluids and reduces gel rigidity, thereby facilitating formulation handling. The resulting SE-doped gel exhibited optimal rheology for rectal administration and for semisolid extrusion (SSE) 3D printing an oral dosage form, consisting of a gastro-resistant shell (33) and a mAb-loaded core with release triggered by intestinal fluids. The therapeutic potential was demonstrated in vivo, where rectal and oral IFX-loaded LMPs significantly ameliorated disease manifestations in a DSS-induced colitis model in rats. Oral biotherapeutic delivery has been widely explored to minimize the adverse effects of systemic treatments. Recent strategies rely on nanocarriers formulated as solutions, administered either as liquid formulations (21,22) or encapsulated into enteric capsules (34). However, these approaches remain limited by formulation stability and challenges in clinical translation. Therefore, to streamline manufacturing while preserving mAb stability, we propose a biocompatible, mucoadhesive gel-based platform for colon-targeted IFX delivery that minimizes systemic exposure and offers a versatile, patient-centric, and translational approach for biotherapeutics.

## Results

### Formulation development and physicochemical characterization

To reduce IFX-related adverse effects while maximizing local therapeutic efficacy, we developed and characterized LMP-based rectal and oral formulations, designed to address different disease localizations and delivery constraints. These systems were tailored to enable controlled and localized mAb release directly at the inflamed intestinal mucosa, either via rectal (InflixiGel; for proctitis) or oral (InflixiPrint and InflixiCaps; for pancolitis) administration. Oral dosage forms can enable colon-specific drug delivery by using coatings responsive to specific gastrointestinal (GI) conditions. InflixiPrint has an LMP-based gastro-resistant layer that dissolves in the intestine, allowing the IFX-loaded LMP in the core to swell and erode, triggering mAb release. This gastro-resistant shell was composed of the phospholipid S80 and vitamin E (S80/VitE_LMP; Fig.1A), whose composition and printing parameters were previously validated (33). The core consisted of the lipid MLO and the hydration-modulating agent SE (MLO/SE_LMP; Fig.1B), and its optimal composition for printability was determined by varying the MLO-to-SE ratio (Fig.1C). Hydration was set at 50% w/w for all MLO/SE LMPs, except for the formulation without SE, where a maximum hydration level was reached with 40% w/w of water (35).

**Figure 1.**
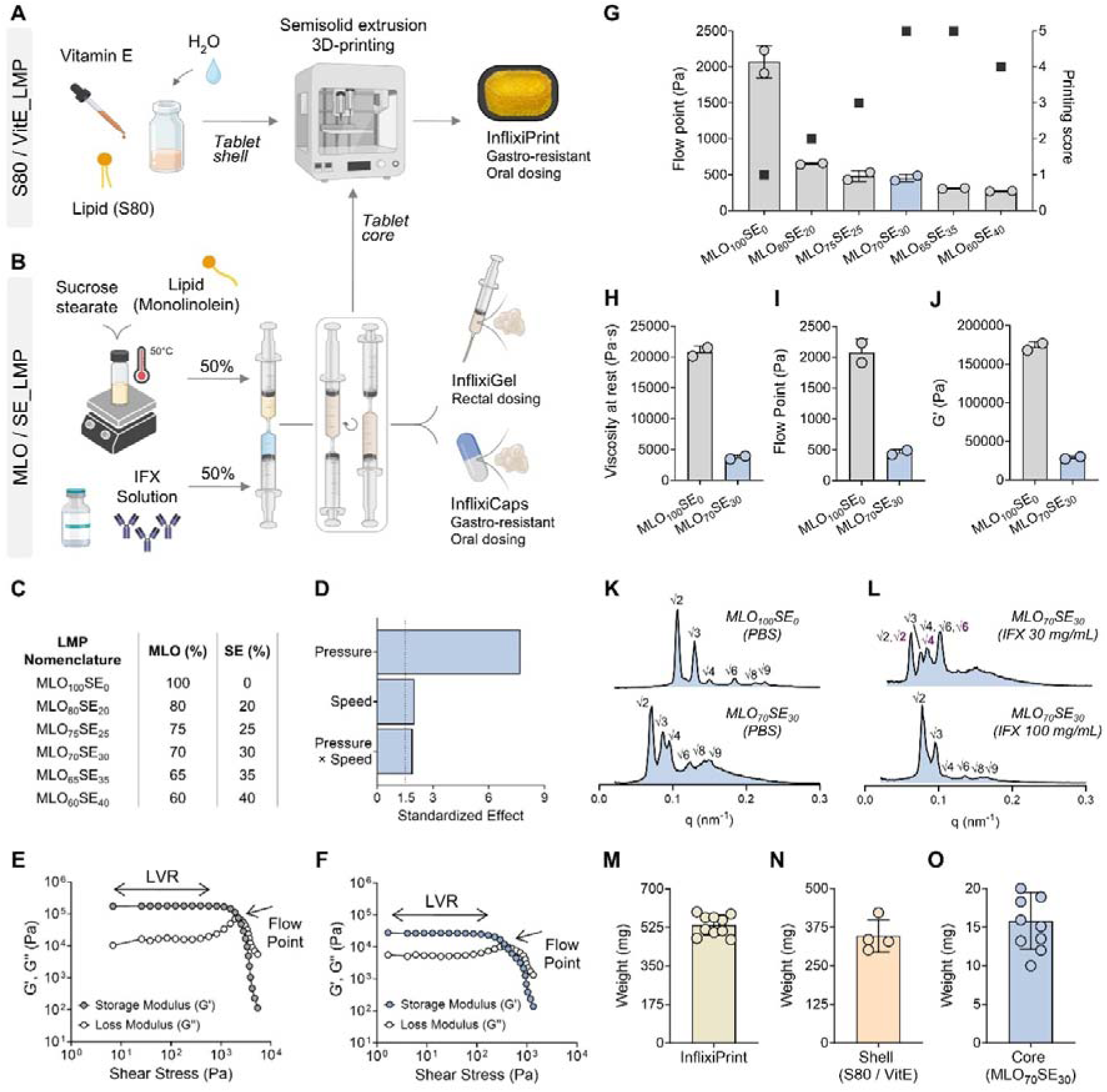
Production and physicochemical characterization of LMPs. (A) Development of S80/VitE_LMP and production of InflixiPrint. (B) Development of MLO/SE_LMP and production of InflixiGel and InflixiCaps. (C) Range of monolinolein (MLO) and sucrose stearate (SE) tested for 3D printing MLO/SE_LMPs. (D) Pareto chart of MLO_70_SE_30_. Amplitude sweeps of (E) MLO_100_SE_0_ and (F) MLO_70_SE_30_. Linear viscoelastic region (LVR) and flow points are indicated by arrows. (G) Correlation between flow point (bars) and maximum printing score (squares) for LMPs formulated with lipid mixtures containing 0-40% SE. (H) Viscosity at rest, (I) flow point, and (J) G’ at the LVR of MLO_100_SE_0_ and MLO_70_SE_30_. (K) Phase identity of MLO_100_SE_0_ and MLO_70_SE_30_ hydrated with PBS and (L) phase identity of MLO_70_SE_30_ hydrated with IFX solution at 30 mg/mL and 100 mg/mL. The Bragg reflections for the cubic Pn3m and Im3m phases are reported above each peak in black and purple, respectively. Average weights of (M) InflixiPrint, (N) S80/VitE shell, and (O) MLO_70_SE_30_ core.

Given the high viscosity and heat sensitivity of lipids and IFX, we employed SSE 3D printing to produce oral dosage forms. The optimal printing parameters for each ink composition were identified using a full factorial Design of Experiment (DoE; Table S1). For each run, a printability score (1 = lowest, 5 = highest) was assigned (Fig.S1), with these values being used to model the influence of the experimental factors and their interactions on printability. All main effects and interactions were evaluated. Factors that did not show statistically significant effects (p≤0.10) were excluded from the final model. The formulation without SE exhibited poor printability, with the ink successfully extruded in only three of the 36 runs (Table S1). Consequently, the data were insufficient for analysis, and the DoE results for this formulation were not reported. The main effects and interaction plots for the remaining formulations are shown in Figure S2, while the Pareto charts show the magnitude of each effect on printability (Fig.1D for MLO_70_SE_30,_ and Fig.S3 for other SE percentages). Pressure had the largest main effect on the printability score across all formulations. Moreover, printing trials showed that higher sugar content reduced the pressure required for ink extrusion. Thus, to further elucidate the rheological behavior of the inks, viscosity-shear rate curves and amplitude sweep tests were performed; the latter was used to determine the storage modulus (G′) within the linear viscoelastic region (LVR) and flow points (Fig.1E, F). Figure 1G shows the correlation between the flow point and the highest printability score obtained for each formulation. The highest printability score (5) was obtained for MLO_70_SE_30_ and MLO_65_SE_35_ (Fig.1G); however, MLO_70_SE_30_ was selected because fewer factors needed to be optimized for printing, as shown in Figure S2. The flow point of MLO_100_SE_0_ was 4.4-fold higher than the average of the other formulations (Fig.1G), explaining its poor printability and highlighting the importance of SE in facilitating extrusion. As shown in Figures 1H-J, the viscosity at rest, flow point and G’ for MLO_100_SE_0_ were 5.6-fold, 3.9-fold and 5.9-fold higher than those of the lead candidate MLO_70_SE_30_, respectively, further supporting this observation.

The influence of SE and IFX on the phase identity of MLO-based LMPs was investigated using SAXS. The first spectrum in Figure 1K (MLO_100_SE_0_) shows Bragg peaks with positions in the ratio √2:√3:√4:√6:√8:√9, characteristic of Pn3m inverse bicontinuous cubic phase. The addition of 30% SE in the lipid phase did not alter mesophase identity (Fig.1K, second spectrum). However, the shift of Bragg peaks to lower q values indicate increased lattice parameter and larger water channels. The swelling effect was further confirmed by the greater water uptake capacity of formulations containing SE (Fig.S4). SE percentages above 40% led to amorphous gels with unknown crystalline domains (Fig.S5). Hydration of MLO_70_SE_30_ with a 30 mg/mL IFX solution shifted the phase boundaries of the system, resulting in the coexistence of Pn3m and Im3m symmetries and a further shift of the Bragg peaks to lower q values (Fig.1L, first spectrum). At a higher IFX concentration (100 mg/mL), the mesophase shrank, reducing its lattice parameter and water channel dimensions (Fig.1L, second spectrum).

The optimal printing parameters identified by the DoE for MLO_70_SE_30_ were a pressure of 110 kPa, a printing speed of 3 mm/s, and an extrusion temperature of 35°C, resulting in 3D-printed tablets of reproducible size and weight (Fig.1M-O, Fig.S6A). For comparison with the 3D-printed tablets (InflixiPrint; Fig.1A), capsules were filled with MLO_70_SE_30_ and coated with Eudragit^®^ FS30D to promote colon targeting (InflixiCaps; Fig.1B). The LMP gel was hydrated with 100 mg/mL IFX solution, delivering 7 mg of IFX per tablet/capsule. The rectal formulation (InflixiGel; Fig.1B) was hydrated with 30 mg/mL IFX solution and delivered at a dose of 1.5 mg IFX per 100 mg of gel. The secondary structure of IFX in the solutions used to hydrate the gels was confirmed using circular dichroism (Fig.S7).

Collectively, these results demonstrate that formulation architecture, manufacturing strategy (SSE 3D printing vs. encapsulation and coating), and compositional tuning via SE incorporation modulate rheological behavior, mesophase structure, processability, and suitability for localized IFX delivery.

### IFX is efficiently encapsulated and released from InflixiGel, InflixiPrint and InflixiCaps

IFX was efficiently encapsulated within the LMP-based formulations, each exhibiting a formulation-specific release profile. The gastro-resistance of InflixiPrint was confirmed by the preservation of the tablet integrity and the absence of detectable IFX or degradation products after incubation under acidic conditions (Fig.2A). Upon transfer to PBS, the gastro-resistant shell of InflixiPrint remained largely intact for up to 75 min, preventing core exposure and resulting in no detectable IFX release after 6 h (Fig.2A, B). In contrast, shell disintegration occurred more rapidly in biorelevant intestinal fluids. In fasted-state simulated intestinal fluid (FaSSIF), near-complete shell disintegration occurred within 45 min, with partial core erosion observed at 75 min. In fed-state simulated intestinal fluid (FeSSIF), both shell and core disintegrated faster, leading to complete tablet disintegration within 75 min (Fig.2A). Accordingly, IFX release after 6 h was higher in FeSSIF (67%) than in FaSSIF (29%) (Fig.2B).

**Figure 2.**
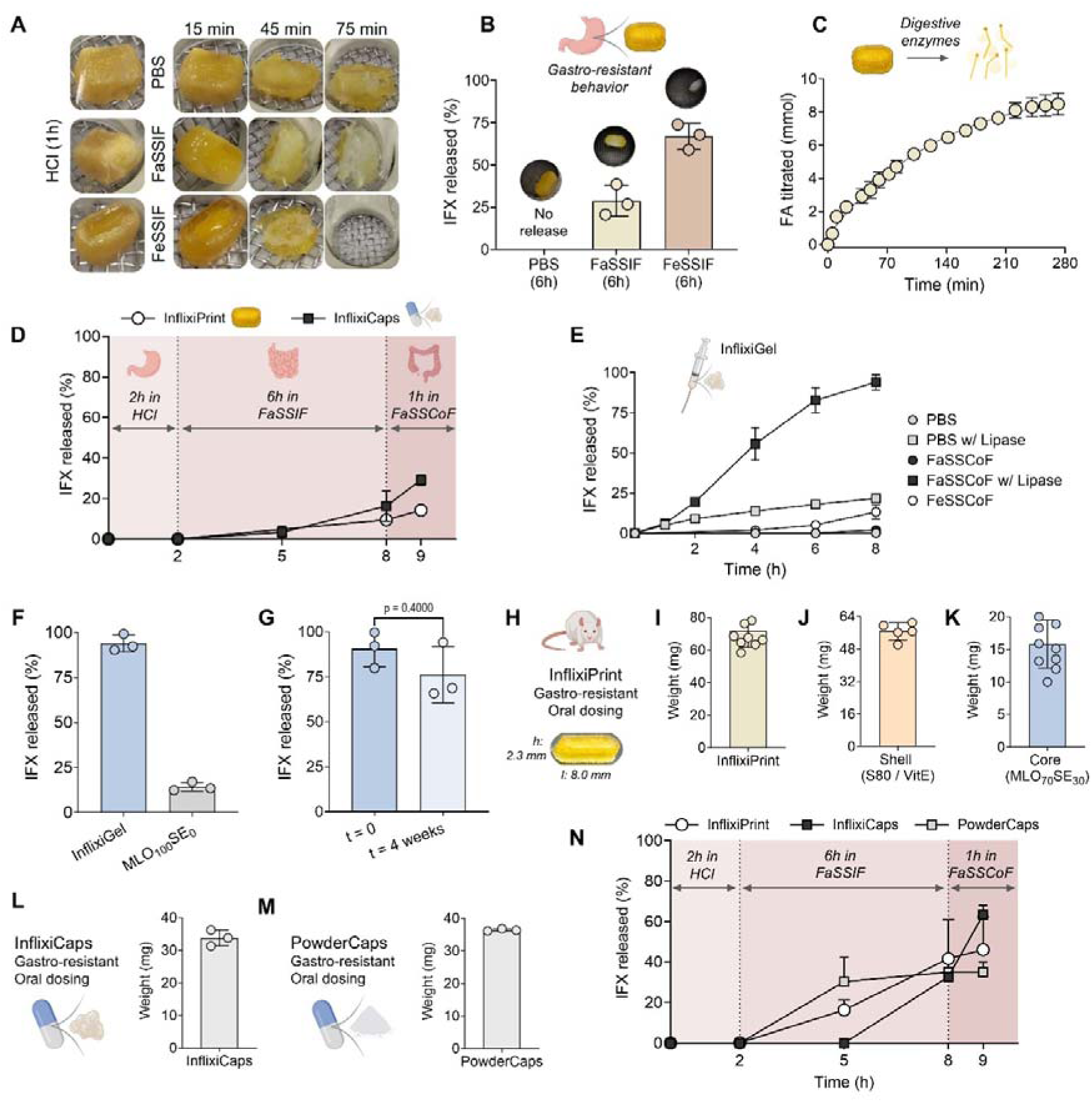
Behavior of formulations in simulated digestive fluids. (A) Disintegration of InflixiPrint in HCl followed by PBS, FaSSIF or FeSSIF. (B) Dissolution of InflixiPrint in PBS, FaSSIF, and FeSSIF. (C) Free fatty acids (FA) originating from InflixiPrint digestion as a function of time (n=2). (D) IFX release from InflixiPrint and InflixiCaps as a function of time in HCl, FaSSIF, and FaSSCoF (n=3). (E) IFX release from InflixiGel in PBS and simulated colonic fluids with or without lipase (n=3). (F) 8-hour IFX release from InflixiGel or MLO_100_SE_0_ in FaSSCoF enriched with lipase. (G) 8-hour IFX release in FaSSCoF enriched with lipase from InflixiGel freshly prepared (t=0) and after storage (t=4 weeks). Miniaturized InflixiPrint (H) dimensions and (I) average weight, and average weight of its (J) shell and (K) core. Average weights of miniaturized (L) InflixiCaps and (M) PowderCaps. (N) IFX release from miniaturized oral dosage forms in HCl, FaSSIF, and FaSSCoF (n=3). Statistical significance in G was evaluated by nonparametric Mann-Whitney test. Adjusted p-values are reported above the comparison.

Lipolysis studies under fasted-state conditions showed progressive digestion of InflixiPrint, with approximately 8.5 mmol of fatty acids (FA) liberated over 240 min (Fig.2C). When the shell and core components were evaluated separately, lipolysis was accelerated, reaching completion within 90 min and releasing 6.5 mmol and 3.2 mmol of FA from the shell and core, respectively (Fig.S8). While these results qualitatively support the disintegration and dissolution data, direct comparison between experiments is not appropriate due to differences in experimental conditions and the absence of digestive enzymes in the disintegration and dissolution assays. The release profiles of InflixiPrint and enteric-coated capsules (InflixiCaps) were compared (Fig.2D), and gastro-resistance was confirmed for both formulations (Fig.S9). In FaSSIF, InflixiPrint and InflixiCaps exhibited similar release profiles, releasing 9.5% and 16% of IFX, respectively, after 6 h. However, in fasted-state simulated colonic fluid (FaSSCoF), IFX release from InflixiCaps was faster (29%) than from InflixiPrint (14%).

The release of IFX from InflixiGel was strongly medium dependent. Drug release was evaluated in PBS, FaSSCoF and fed-state simulated colonic fluid (FeSSCoF), both with and without lipase (5000 U/mL) to mimic the presence of this enzyme in the inflamed colonic mucosa (36–38) (Fig.2E). The rectal dosage form consisted solely of MLO_70_SE_30_ and, despite the absence of the S80/VitE shell, no IFX release was detected over 8 h in PBS. In simulated colonic fluids, IFX release remained limited (2% in FaSSCoF and 13% in FeSSCoF), with higher release in FeSSCoF attributed to its greater bile salt content. Lipase addition significantly increased IFX release in PBS and FaSSCoF (22% and 94%, respectively), indicating that release is primarily driven by enzymatic erosion of the LMP rather than passive diffusion through aqueous channels. Gel erosion and drug release were not observed in lipase-enriched FeSSCoF likely due to enzyme inactivation by glucose present in this medium but absent in FaSSCoF (39).

To further investigate the role of the swelling agent, we compared IFX release from InflixiGel (MLO_70_SE_30_) and from a gel without SE (MLO_100_SE_0_). InflixiGel released nearly 100% IFX, whereas the release from MLO_100_SE_0_ was limited to 14% (Fig.2F), confirming the critical role of the SE in promoting drug release. This feature can be explained by the high hydrophilic-lipophilic balance (HLB) value of the SE, which functions as a solubilizing agent for the LMP and facilitates its erosion. Formulation stability was assessed by comparing IFX release from freshly prepared InflixiGel with the release from gels stored at 4°C for four weeks. Comparable drug release amounts demonstrate preserved formulation stability (Fig.2G).

For in vivo evaluation, the InflixiPrint design was scaled down to match the dimensions of rat capsules (Torpac, size 9) (Fig.2H, Fig.S6B) The weights of the core, shell, and total tablet were consistent across replicates (Fig.2I-K). For comparison, enteric-coated capsules were filled with the same gel used to print the core of the 3D-printed tablet (InflixiCaps; Fig.2L). We also investigated whether the LMP matrix was necessary for the oral delivery of IFX by filling colon targeting capsules with a wet powder mixture of mannitol and IFX solution (PowderCaps; Fig.2M). Despite their different compositions, InflixiCaps and PowderCaps exhibited comparable weights (Fig.2L, M). Miniaturized oral dosage forms were designed to deliver a dose of 5 mg/kg.

IFX release from the rat-sized oral dosage forms was assessed under gastric and fasted intestinal conditions and compared with the release profile from human-sized printlets and capsules (Fig.2N). Gastro-resistance was confirmed by the absence of detectable IFX or degradation products after incubation in acidic conditions. For InflixiPrint, the change in medium composition from FaSSIF to FaSSCoF did not markedly affect the release rate, which is consistent with the observations for the human-sized printlet. In contrast, InflixiCaps exhibited a burst release upon transfer to FaSSCoF, releasing approximately 30% of IFX within 1 h. PowderCaps exhibited faster release in FaSSIF, releasing 30% IFX within 3 h; however, no further release occurred in FaSSCoF. Although the coating of the capsules did not remain intact for the full 6 h in FaSSIF, its thickness was limited to ensure capsule compatibility with the gavage applicator. The maximum number of coating layers was applied while maintaining suitability for in vivo administration.

### Gastrointestinal transit of InflixiCaps and PowderCaps and colonic retention of InflixiGel

A pilot study was conducted to evaluate the feasibility of oral gavage of InflixiPrint and capsule formulations in rats. Although the 3D-printed dosage form was sufficiently robust for human handling, its lipid nature caused adhesion to the small gavage applicator and to the animals’ esophagus. This made administration difficult and uncomfortable, at least in small rodents like rats. Therefore, to prioritize animal welfare and to fulfil the 3R principles, this formulation was excluded from in vivo testing, and the GI transit was evaluated for InflixiCaps and PowderCaps. Following oral gavage of barium sulfate-loaded capsules, computed tomography (CT) imaging of the anesthetized rats was performed at predefined time points. Imaging at t=0 confirmed gastric localization of both formulations immediately after administration (Fig.3A, top). At 4 h post-gavage, InflixiCaps had progressed to the cecum, and they were consistently detected in the colon by 6 h (Fig.3A, left). In contrast, PowderCaps remained in the stomach at both 4 and 6 h (Fig.3A, right). These observations were reproducible across all animals (n=3/group; Fig.3B), demonstrating that the LMP matrix facilitated efficient GI transit and timely colonic arrival.

**Figure 3.**
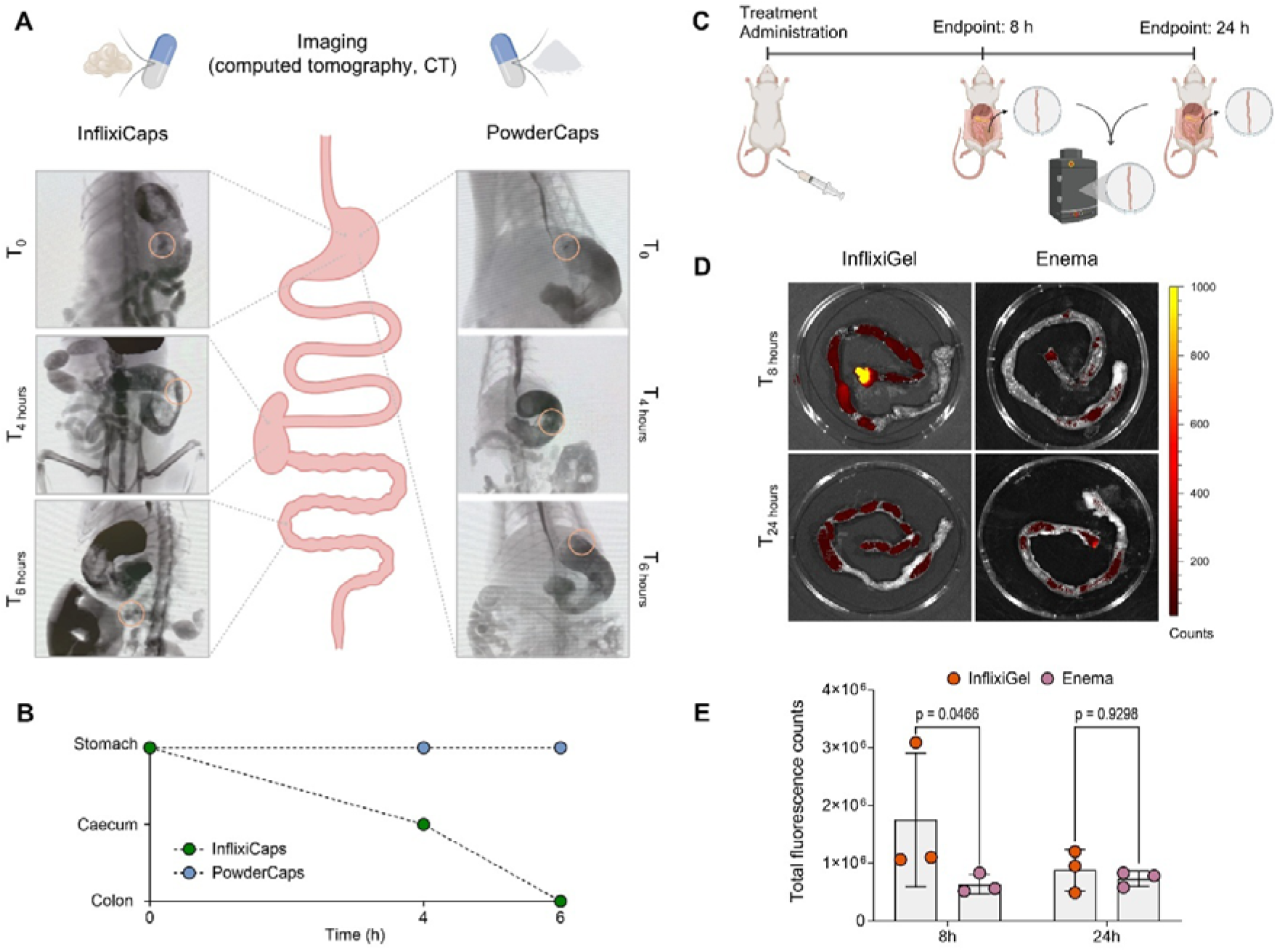
GI transit of oral formulations and colonic retention of rectal formulations. (A) Representative CT images of the rat GI tract over time after gavage of InflixiCaps and PowderCaps. (B) Location of InflixiCaps and PowderCaps in the GI tract over time (n=3). (C) Experimental setup for assessing colonic mucoadhesion of rectal formulations. (D) Representative images of colons and (E) total fluorescence counts at 8 and 24 h after administration of rectal formulations (n=3). Statistical significance was evaluated by two-way ANOVA with Tukey’s post hoc analysis. Adjusted p-values are reported above the comparison.

Insufficient mucoadhesion and leakage can compromise the efficacy of rectal formulations; therefore, we evaluated the colonic residence time of InflixiGel compared with an enema (aqueous IFX solution formulated at the same concentration as in the gel). Fluorescently labelled rectal formulations were administered, and the rat colons were harvested 8 and 24 h post-administration for fluorescence intensity analysis (Fig.3C). Representative images are shown in Figure 3D. After 8 h, the fluorescence signal in colons from InflixiGel-treated rats was higher than that in the enema group, indicating stronger mucosal adhesion. After 24 h, the fluorescence signal in the InflixiGel-treated group decreased approximately two-fold and reached values comparable to those observed in the enema group (Fig.3E). Notably, the signal of the gel-treated colons was higher than that of the untreated controls (Fig.S10), confirming the sustained colonic presence of InflixiGel.

### IFX Systemic Absorption and Colonic Retention

Systemic absorption and colonic retention of IFX were first characterized in DSS-treated rats following subcutaneous administration (2 mg/kg) as a reference for our localized delivery systems. IFX reached a C_max_ of 23 µg/mL at 4 days post-injection, with an AUC of 230 µg/mL×d over 22 days (Fig.4A-C). Colonic IFX levels increased 3.8-fold by day 7 (0.16 µg/g tissue) and remained stable thereafter (Fig.4D).

**Figure 4.**
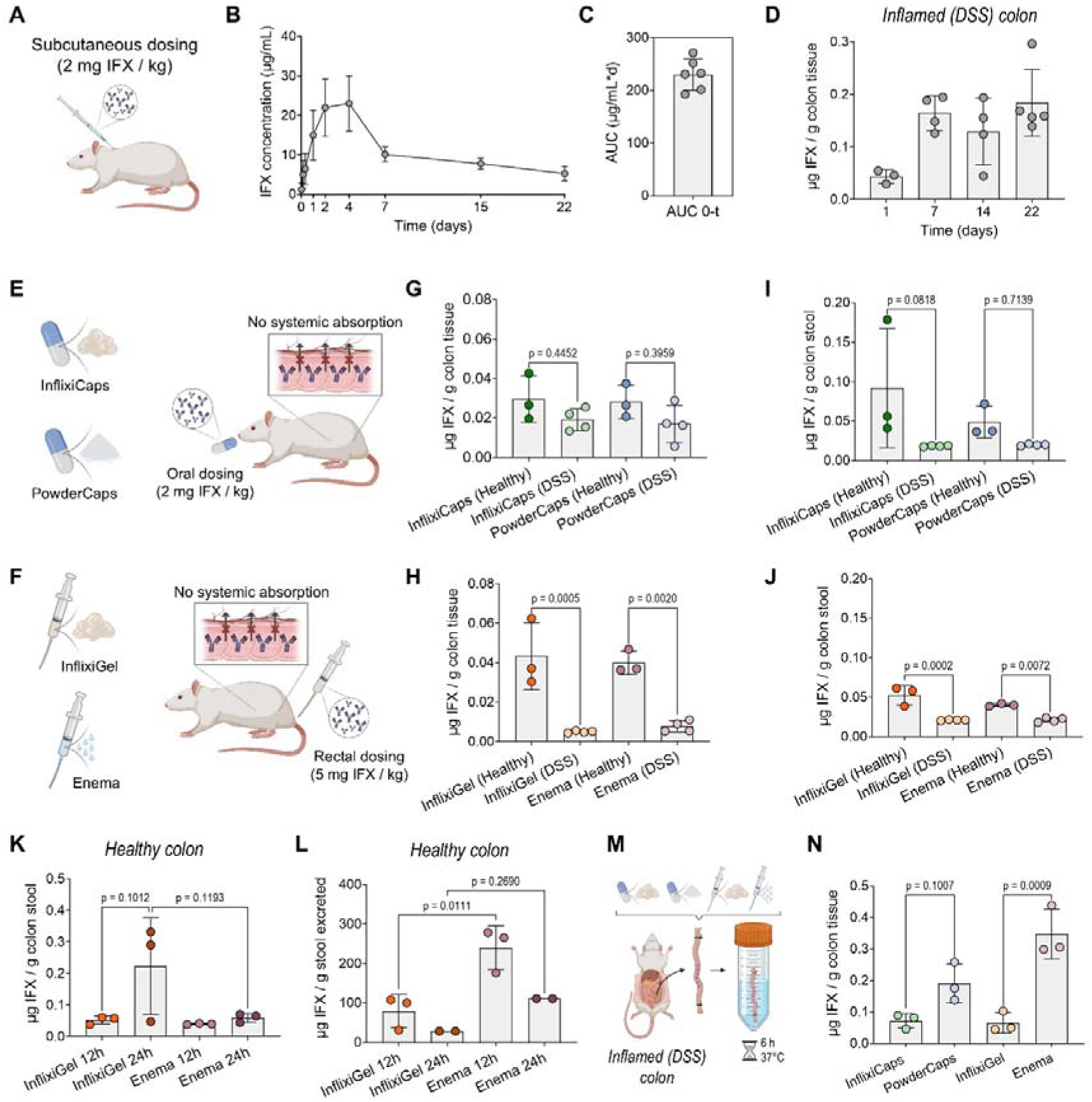
Blood and tissue pharmacokinetics after IFX subcutaneous, oral and rectal administrations. (A) Subcutaneous IFX injection to DSS-treated rats. (B) Serum concentration of IFX over time (n=6), (C) total systemic exposure (AUC) and (D) colonic retention after subcutaneous injection. IFX was not detected in the serum of DSS-treated rats, neither after (E) oral nor (F) rectal administration. Colonic concentration of IFX in healthy and DSS-treated rats after (G) oral and (H) rectal administration. Concentration of IFX in the colonic stools of healthy and DSS-treated rats after (I) oral and (J) rectal administrations. Concentration of IFX in (K) colonic stools and (L) excreted stools of healthy rats after rectal administration. (M) Ex vivo experimental setup and (N) IFX tissue concentration after 6-hour incubation of oral and rectal formulations. Statistical significance was evaluated by one-way ANOVA with Tukey’s post hoc analysis. Adjusted p-values are reported above the comparison.

The systemic absorption of mAbs is associated with adverse effects (40). To minimize these risks, our formulations were designed for localized mAb delivery, aiming for high drug concentrations in the inflamed colon and limited systemic exposure. Accordingly, no detectable systemic absorption of IFX was observed following oral (InflixiCaps and PowderCaps, 2 mg/kg) and rectal (InflixiGel and enema, 5 mg/kg) administrations (Fig.4E, F). Whereas the oral dose was selected based on the systemic reference dose, the rectal dose was guided by our previous study, which showed that half of the LMP gel was excreted after administration (24).

Colonic retention was evaluated 12 h post-rectal administration and 24 h post-oral administration in both healthy and DSS-treated rats (Fig.4G, H). After oral administration, colonic IFX levels in healthy rats were slightly higher for both InflixiCaps and PowderCaps, although the differences were not statistically significant (p>0.05). Similarly, rectal administration resulted in higher IFX tissue concentrations in healthy rats compared with DSS-treated counterparts. Stool analysis revealed lower IFX levels in colitic animals (Fig.4I, J). To assess retention over time, colonic tissue and stool samples were collected from healthy rats 12 and 24 h after rectal administration of InflixiGel and enema. IFX tissue concentrations were comparable at both time points for both formulations (Fig.S11). However, higher IFX amounts were detected in the colon contents at 24 h for InflixiGel, suggesting prolonged local retention (Fig.4K). Analysis of excreted stools collected throughout the experimental period further showed higher IFX levels in enema-treated rats (Fig.4L), consistent with its lower mucoadhesion (Fig.3E).

The lower IFX levels detected in colonic tissues of DSS-treated rats might result from rapid binding and consumption of the antibody by TNF-α, which is overexpressed in the inflamed intestinal mucosa of IBD patients (41–43). To test this hypothesis, we harvested colons from rats and assessed the retention of IFX in an ex vivo setup, where formulations were added inside the colon and both ends of the tissue were sealed (Fig.4M). In the ex vivo setup, all formulations led to markedly higher tissue concentrations of IFX compared to in vivo (Fig.4N). Non-LMP formulations (PowderCaps and enema) showed higher IFX tissue retention, highlighting the controlled release profile of LMP-based gels (InflixiCaps and InflixiGel).

The rapid GI transit of InflixiCaps, reaching the colon within 6 h, together with the low IFX levels detected in colonic tissues and stools, supports a daily dosing regimen for efficacy studies. Similarly, IFX concentrations in inflamed colonic tissues and stools following rectal administration, along with the higher excretion observed with the enema, also support a daily dosing frequency of rectal formulations.

### Prophylactic Efficacy of InflixiCaps and InflixiGel in DSS-induced colitis rats

The protective effects of InflixiCaps and InflixiGel against colonic inflammation were investigated using a DSS-induced colitis model in Sprague-Dawley rats. Different to other chemically induced colitis models, DSS models allow modulation of disease severity through concentration and dosing frequency (44). However, inflammatory response is modulated by the intestinal microbiome (45). DSS has been employed at a range of concentrations (4-8%) to induce acute colitis in Sprague-Dawley rats (34,46–48). In this study, DSS administration was standardized by using 4% and 6% DSS for seven days to induce moderate and severe colitis, respectively. In both inflammation severity models, no significant reduction in body weight was observed across all treatment groups (<5% reduction; Fig.S12).

In the moderate colitis model, untreated animals developed soft stools with occasional diarrhea and bleeding, as well as progressively increasing disease activity index (DAI) values, which were similar between untreated males and females (Fig.S13). Both InflixiGel and InflixiCaps produced a modest early reduction in DAI, which became statistically significant from day five onward, resulting in significantly lower DAI values compared with untreated animals (Fig.5A). At day seven, fecal lactoferrin, a validated marker of intestinal inflammation (49–51), was significantly elevated in untreated rats relative to naïve controls (p=0.0353) and significantly reduced by both localized IFX formulations (Fig.5B). Colon length, an indicator of DSS-induced tissue damage, was reduced in untreated animals, particularly in males (Fig.S14). Treatment with InflixiGel and InflixiCaps trended toward preservation of colon length; however, this difference was not statistically significant (p>0.05; Fig.5C). In this model, tissue concentrations of TNF-α were comparable across all experimental groups (Fig.5D).

**Figure 5.**
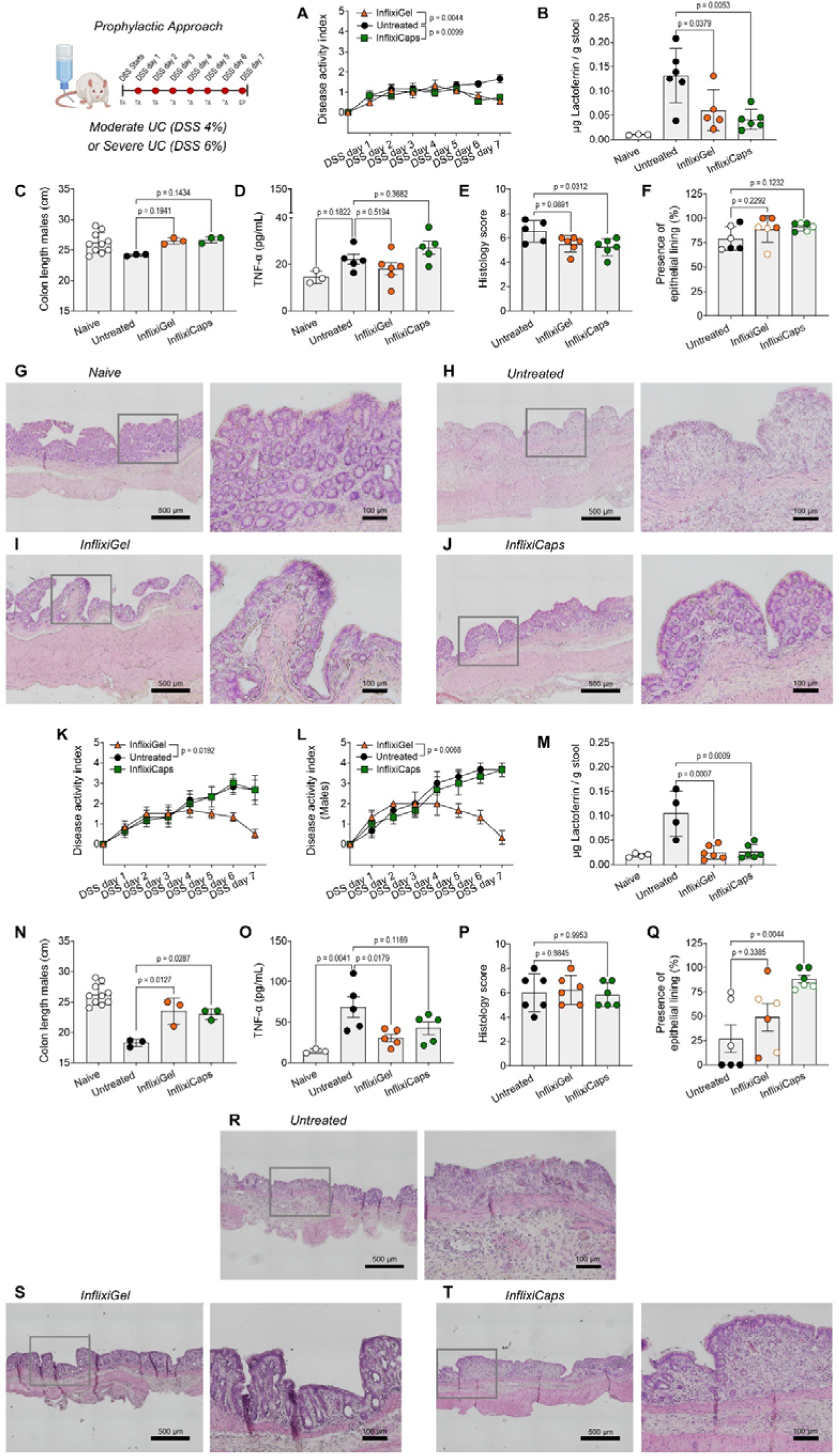
InfixiGel and InflixiCaps prophylactic efficacy in moderate and severe in DSS-exposed rats. Animals received 4% DSS in A-J. (A) Daily disease activity index. (B) Fecal lactoferrin concentration, (C) colon length, and (D) colon TNF-α concentration in naïve, untreated, and treated rats after DSS cycle. (E) Histology score and (F) presence of epithelial lining in males (closed symbols) and females (open symbols) after DSS cycle. Representative histology pictures of (G) naïve, (H) untreated, (I) InflixiGel-treated, and (J) InflixiCaps-treated animals after DSS cycle. Animals received 6% DSS in K-T. Daily disease activity index of (K) female and male rats and (L) male rats. (M) Fecal lactoferrin concentration, (N) colon length, and (O) colon TNF-α concentration in naïve, untreated, and treated rats after DSS cycle. (P) Histology score and (Q) presence of epithelial lining in males (closed symbols) and females (open symbols) after DSS cycle. Representative histology pictures of (R) untreated, (S) InflixiGel-treated, and (T) InflixiCaps-treated animals after DSS cycle. Sample size was n=6 rats/group. Statistical significance was evaluated using Linear Mixed-Effects (A, K, L), and one-way ANOVA with Tukey’s post hoc analysis (B-F, M-Q), for which outliers were identified by Grubbs’ test (α=0.05) and removed in B, D, E, M, O. Adjusted p-values are reported above the comparison. TA: treatment administration, EP: endpoint.

Histologically, InflixiGel and InflixiCaps mitigated epithelial injury, preserved crypt architecture, and maintained goblet cell populations, with reduced lymphocytic infiltration and submucosal edema compared to untreated rats (Fig.5E). In addition, the colonic mucosa was better preserved in the treated groups, although not statistically different than the untreated group (Fig.5F). Compared to naïve animals, untreated DSS-exposed animals exhibited severe mucosal damage characterized by marked crypt distortion, extensive goblet cell depletion, and a fragmented lining epithelium (Fig.5G, H). Light lymphocytic infiltration was observed in both the mucosa and submucosa, together with prominent submucosal edema. InflixiGel-treated rats showed moderate crypt distortion and detectable goblet cells (Fig.5I). Mild lymphocytic infiltration was mainly confined to the mucosa, whereas submucosal edema remained at a similar extent as in the other groups. Similarly, InflixiCaps-treated animals exhibited preserved crypt architecture and maintained goblet cell populations (Fig.5J). Mild lymphocytic infiltration was primarily restricted to the mucosa, and submucosal edema was observed at levels comparable to those in the other treatment groups. Together, these results demonstrate that both strategies for localized IFX delivery provide significant protection against DSS-induced colonic inflammation. In the severe colitis model, untreated animals exhibited markedly higher DAI values than those observed in the moderate colitis model, due to persistent diarrhea and frequent rectal bleeding. In this setting, InflixiCaps did not significantly attenuate DAI, whereas InflixiGel robustly reduced DAI from day five onward (Fig.5K). These results highlight the advantage of rectal administration for achieving high local drug concentrations. Disease severity was sex-dependent in this model, with males developing more severe disease than females (Fig.S15). In males, InflixiGel nearly normalized DAI by day seven (Fig.5L), highlighting the strong efficacy of the rectal formulation under severe inflammatory conditions. Fecal lactoferrin levels were elevated in untreated animals and reduced to near-naïve levels in InflixiGel and InflixiCaps treatment groups (Fig.5M). Untreated males exposed to 6% DSS displayed significantly shorter colons than naïve animals, whereas this effect was less evident in females (Fig.S16). Accordingly, the colon length was analyzed in males (Fig.5N). Both InflixiGel and InflixiCaps preserved colon length relative to untreated controls, with values comparable to those of naïve animals. In contrast to the moderate colitis model, severe colitis was associated with a marked TNF-α component and, under these conditions, both InflixiGel and InflixiCaps significantly reduced tissue TNF-α concentrations (Fig.5O), restoring levels to those observed in naïve controls.

Epithelial damage and immune cell infiltration scores were comparable among groups (Fig.5P), demonstrating that improvements in macroscopic outcomes and biomarkers were not accompanied by corresponding histological healing. However, qualitative histological analysis revealed differences in mucosal integrity among treatment groups. InflixiCaps markedly preserved the epithelial lining compared with untreated animals (Fig.5Q). Particularly, untreated males exhibited complete loss of the epithelial lining. Additionally, the untreated group displayed crypt distortion, goblet-cell depletion, and widespread lymphocytic infiltration (Fig.5R). In contrast, InflixiGel preserved crypt architecture, maintained goblet cells, and restricted lymphocytic infiltration to the mucosa (Fig.5S). InflixiCaps produced a similar protective pattern, with moderate crypt distortion and sporadic goblet-cell preservation (Fig.5T). Importantly, this histological profile closely resembled that observed in the moderate DSS model, indicating that both formulations effectively protect the colonic mucosa under severe inflammatory stress, with formulation-specific advantages: InflixiCaps predominantly maintained the epithelial lining, whereas InflixiGel, likely through topical contact with the mucosa, preserves the crypt architecture.

## Discussion

Despite their clinical success in UC treatment, systemically administered mAbs are associated with dose-limiting toxicity, immunogenicity, and suboptimal drug exposure to the inflamed mucosa. Intravenous (IV) or subcutaneous delivery of IFX results in high systemic exposure despite the disease being confined to the colon. Here, we addressed this fundamental mismatch between drug delivery route and disease localization by developing rectal and oral LMP-based formulations that successfully delivered IFX to the inflamed colon while minimizing systemic absorption. Using complementary in vitro, ex vivo, and in vivo approaches, we show that LMPs protect IFX from gastrointestinal degradation, enable its release in response to colonic biochemical cues, and enhance mucosal retention. Importantly, this localized delivery translated into robust therapeutic efficacy in DSS-induced colitis, as evidenced by reductions in disease activity, inflammatory biomarkers, and histological damage.

### Formulation development and physicochemical characterization

Oral delivery of biologics requires gastro-resistant protection, as these molecules are rapidly degraded in the stomach (52). Our research group previously developed a gastro-resistant 3D-printed tablet using S80/VitE_LMP (33), which informed the selection of this gel as the protective shell for InflixiPrint. This LMP exhibits mixed lamellar and hexagonal phases, both associated with slower diffusion rate compared to inverse bicontinuous cubic phases (23). In addition, the S80/VitE_LMP contains only 20% water, limiting the incorporation of hydrophilic molecules within its aqueous channels. In contrast, cubic mesophases allow for higher water content, increased loading of hydrophilic molecules, and faster drug diffusion (23). However, their narrow water channel diameter can restrict the release of macromolecules (26,27). To overcome this limitation, hydration-modulating additives such as SE have been incorporated into cubic mesophases (28,29). Accordingly, the InflixiPrint core was designed using an MLO-based LMP containing SE at an optimized concentration to enable successful 3D printing while maintaining mechanical stability. Increasing SE content reduced the flow point and improved extrudability. The structural effect of SE on LMPs has been reported (28), and its influence on rheological behavior has been linked to a reduction in apparent viscosity (53), consistent with our findings. Moreover, a direct link between water channel dimensions and structural strength has already been reported, which is in line with our observation (54).

SAXS measurements were conducted on the LMP selected for printing the core (MLO_70_SE_30_). A previous study reported significant swelling effect by adding SE to the lipid mixture, accompanied by a phase transition from Pn3m to Im3m (28). The highly hydrophilic headgroup of SE decreases the critical packing parameter (CPP) and reduces the curvature of the lipid bilayer towards the aqueous phase, resulting in phase transition and enlarged water channels. However, the curvature of the lipid bilayer is determined not only by the molecular structure of its constituent lipids and additives, but also by external factors such as salts present in the aqueous phase (30,55,56). Here, the addition of 30% SE induced only minor swelling when hydrated with PBS, and a complete phase transition to Im3m was not observed. Negrini et al. (28) hydrated the mesophase using HEPES buffer, which has weaker charge density compared with PBS. Chaotropic ions, characterized by weaker hydration and lower charge density, have been reported to increase mesophase hydration and the lattice parameter. In contrast, kosmotropic salts, such as Na_2_HPO_4_ in PBS, decrease the lattice parameter by strengthening ion-water interactions, leading to an increased bilayer curvature (57). These differences explain the overall reduced swelling capacity of our system. Hydration of the mesophase with a 30 mg/mL IFX solution further increased the water channel diameter, and SAXS data revealed a coexistence of Pn3m and Im3m phases under these conditions. We hypothesized that the accommodation of large antibody molecules within the water domains caused frustration in bilayer packing, triggering the transition from Pn3m to Im3m. Conversely, hydration of the mesophase with a highly concentrated IFX solution (100 mg/mL) reduced the water channel diameter. This concentrated solution contained not only more IFX but also a higher level of phosphate salts from the lyophilized powder (Na_2_HPO_4_ and Na_₂_HPO_₄_), which promoted water channel shrinkage (57).

### IFX is efficiently encapsulated and released from InflixiGel, InflixiPrint and InflixiCaps

Lipid-based solid oral dosage forms have demonstrated clinical and commercial success (58,59). However, their formulation remains challenging because of the high viscosity and heat sensitivity of many lipids, which complicate conventional tableting processes (33,60). Consequently, most existing lipid-based solid oral formulations rely on encapsulation, often using gelatine capsules (59). SSE 3D printing provides an alternative approach for producing lipid-rich solid oral dosage forms. The printable S80/VitE_LMP developed by our group has an 80% lipid content. This formulation was shown to be gastro-resistant and to undergo digestion in simulated intestinal fluids, forming colloidal structures that enhance the solubility of poorly water-soluble drugs (33).

In this study, InflixiPrint also maintained its integrity in acidic conditions, confirming gastro-resistance. Disintegration and subsequent dissolution are critical determinants of oral drug absorption and can represent a rate-limiting step in the absorption process (61). While lipid-based solid oral dosage forms typically include disintegrants and/or surfactants to enable these processes, InflixiPrint was deliberately designed for slow disintegration in intestinal fluids to achieve delayed, colonic drug release. Slower disintegration and dissolution were observed in FaSSIF, which is consistent with its lower bile salt concentration compared to FeSSIF. Therefore, administration under fasted conditions is expected to prolong disintegration and favor a delayed release. Considering that the average small intestinal transit time is approximately 3-4 h (62), the <30% drug release within 6 h suggests that a substantial fraction of IFX remains in the core until it reaches the colon, where complete release can occur. IFX release from InflixiPrint was compared with InflixiCaps under fasted conditions. Both formulations exhibited comparable release profiles; however, InflixiCaps showed higher IFX release in FaSSCoF. This difference likely arises from the pH-responsive coating on InflixiCaps, which dissolves above pH 7, facilitating a faster release. In contrast, InflixiPrint demonstrated a more gradual shell disintegration, resulting in a slower release profile.

InflixiGel demonstrated a high yet gradual IFX release (up to 8 h) in FaSSCoF enriched with lipase, an enzyme present in the inflamed colonic mucosa (36–38). These findings indicate that IFX release is enhanced under inflammatory conditions, potentially increasing mAb concentration in inflamed sites. The limited IFX release from the LMP lacking SE (MLO_100_SE_0_), contrary to what was observed for InflixiGel, likely results from lipase-mediated hydrolysis of the sugar ester (63–65), and confirms the critical role of SE in enabling inflammation-triggered release from InflixiGel in the colon. Interestingly, the lipase-mediated release was less pronounced in PBS, suggesting a synergistic effect of lipase and FaSSCoF in promoting drug release. This highlights the importance of evaluating release kinetics under physiologically relevant conditions to accurately predict the in vivo performance.

Stability studies on mAb aqueous solutions have evaluated the effects of excipients, pH, temperature, and other variables (66–68), as well as the impact of their incorporation into delivery systems (69–71). Here, we demonstrated that MLO/SE_LMPs enable mAb encapsulation at high concentrations (30 mg/mL) and effectively prevent protein aggregation and degradation during storge, as evidenced by comparable amounts of active IFX released from the gel after storage. We hypothesized that the 3D mesophase structure is partially responsible for preserving the mAb monomeric form and, therefore, its biological activity. Indeed, LMPs have been widely employed for membrane protein crystallization (72,73), supporting our hypothesis that IFX is comfortably accommodated within the LMP aqueous channel network. Notably, additional stability studies are required to assess the long-term stability of these formulations. We recently validated the dual-syringe method for LMP production, enabling rapid mixing of the aqueous and lipid phases by the patient (74). This approach allows administration of freshly prepared gels and overcomes potential long-term stability limitations. Additionally, for oral dosage forms, 3D printing enables cost-efficient, personalized manufacturing and supports on-demand production. By bringing manufacturing closer to the patient, this strategy may facilitate the use of biotherapeutics with limited stability.

Scaling down the formulation size for preclinical studies resulted in only minor changes in formulation behavior. The slightly faster release observed in FaSSIF for the rat-sized printlet, compared to the human-sized one, may be attributed to its thinner shell (0.2 mm vs. 1 mm; Fig.S6), which is more rapidly digested in simulated intestinal fluids, resulting in earlier exposure of the IFX-loaded core. The faster release from PowderCaps compared to InflixiCaps, despite the same coating, is likely due to the absence of the LMP matrix as IFX is readily available once the PowderCaps coating dissolves, whereas, in InflixiCaps, IFX diffusion is slowed by its encapsulation within the LMP.

Rectal and oral delivery of biotherapeutics have been widely investigated; however, no commercial dosage form currently enables localized mAb delivery to the inflamed colon. Conventional rectal formulations show poor retention and insufficient drug absorption, with drug-excipient interactions potentially impairing mAb stability. Enteric capsules filled with protein-based solutions and standard tableting procedures are also unsuitable, as they can induce protein degradation. Here, we propose LMP-based formulations that directly immobilize the antibody within a protective gel matrix. This approach enhances IFX stability, simplifies manufacturing and enables controlled colonic release through matrix erosion rather than rapid pH-triggered dissolution. Moreover, unlike nanoparticle-based systems (such as liposomes, micelles, and lipidic nanoparticles), LMP gels are not internalized by infiltrating macrophages following oral or rectal administration.

### Gastrointestinal transit of InflixiCaps and PowderCaps and colonic retention of InflixiGel

The high viscosity and stiffness of LMPs can limit their use as a standalone delivery system (75,76). We demonstrated that tuning the mesophase composition allows SSE 3D printing of solid oral dosage forms composed exclusively of LMPs. However, the difficulty of administering these systems to animals restricts in vivo testing. Other studies have encountered similar challenges when delivering LMPs orally and have addressed this by administering a mesophase precursor instead (76,77). A previous study showed that cubic mesophases formed in-situ after oral administration of lipids containing a poorly water-soluble drug, resulting in sustained drug release (77). Another strategy, the same adopted by us, is using enteric capsules filled with LMPs (78). However, gastric emptying is governed by a complex interplay between physiological and formulation-related factors and often represents a rate-limiting step in the absorption of orally administered drugs. Inconsistencies in the gastric emptying of size 9 capsules with comparable average weights have been documented in anesthetized rats (79). To minimize anesthesia-induced effects on GI motility, our study incorporated a 4 h interval between measurements, as motility impairment has been reported for up to 2 h after isoflurane exposure (80).

Hydration and swelling of Eudragit^®^ FS30D films in gastric fluids have been previously reported (81). This behavior potentially allowed water to diffuse into the capsules, thereby hydrating their cores while still preventing drug release. Consequently, mannitol in PowderCaps and LMP in InflixiCaps were hydrated. Owing to its greater swelling capacity, LMP absorbs more water than powder alone (mannitol), resulting in increased weight. We further characterized this behavior in vitro by incubating InflixiCaps and PowderCaps in water. Whereas PowderCaps remained floating at the surface, InflixiCaps sank, providing a possible mechanistic explanation for the faster gastric emptying observed for InflixiCaps (Fig.S17).

Previously, MLO-based LMPs demonstrated superior colonic retention in mice compared to enemas (24). Our findings confirm that incorporation of SE further enhances mucoadhesive behavior. This effect is likely mediated by weak secondary interactions between the hydrophobic backbone of sugar esters and mucin strands (82), promoting prolonged mucosal adhesion.

The pharmacokinetic profile observed following subcutaneous injection was consistent with previously reported data for anti-TNF antibodies in rodents (83,84) and humans (85). Elevated colonic levels of IFX in inflamed tissues have been attributed to TNF-α overproduction and the consequent recruitment of anti-TNF agents, allowing inflamed tissues to act as drug reservoirs (86). However, under severe inflammatory conditions, reduced tissue levels of anti-TNF agents have been reported, likely due to accelerated clearance (87). We hypothesized that the lower IFX concentrations detected in colonic tissues and stools of DSS-treated rats following oral and rectal administrations result from its immediate consumption by TNF-α in the colonic mucosa. This hypothesis was supported by ex vivo experiments using inflamed colonic tissues, where higher IFX retention was observed. TNF-α levels in the tissue rapidly decrease after harvesting, explaining the increased colonic IFX concentrations observed ex vivo. These findings demonstrate that IFX remains structurally stable and effectively penetrates inflamed colonic tissues. Additionally, the controlled-release properties of the LMP-based systems were evidenced by the lower IFX colonic retention from InflixiCaps and InflixiGel compared with the non-LMP formulations. However, the experiment was conducted in a closed system with both ends of the colon sealed, which enhances formulation-tissue contact and promotes maximal retention. Particularly, the high IFX concentrations in the enema-treated colon are not representative of in vivo conditions, where this formulation leaks rapidly after administration and shows poor colonic retention.

Colon-targeted IFX delivery was successfully achieved using LMP-based formulations. The oral formulation, InflixiCaps, resisted GI transit and reached the colon within 6 h, where degradation of the coating exposed the LMP core and enabled local drug release. In parallel, the rectal formulation, InflixiGel, remained strongly adhered to the colonic mucosa for more than 8 h. Together, these two delivery platforms enabled IFX release directly at inflamed sites, thereby avoiding the adverse effects associated with systemic and infusion-based anti-TNF therapy.

### Prophylactic Efficacy of InflixiCaps and InflixiGel in DSS-induced colitis rats

Patients who responded to anti-TNF agents for induction of remission, are recommended to continue treatment with the same drug for maintenance of remission (6). In clinical practice, IFX is often the biologic of choice; however, maintenance therapy requires repeated IV infusions every eight weeks (88), placing a substantial burden on both patients and healthcare systems. While subcutaneous administration offers a more convenient, at-home option compared with IV infusion, it is associated with a similar anti-drug antibody development profile (89), which can lead to impaired therapeutic response and adverse effects (90). It has been reported that IFX concentrations during maintenance phase exert the strongest influence on the risk of anti-drug antibody development (91). In this context, localized therapeutic approaches that reduce systemic exposure and immunogenicity are particularly important during maintenance therapy. Our oral and rectal LMP-based formulations may provide a convenient and needle-free alternative for patients while also reducing the burden on the healthcare system.

The protective effects of the LMP-based formulations were evaluated in DSS-induced models of moderate and severe colitis. Regardless of inflammation severity, body weight could not be used as a reliable readout of disease activity as it remained stable or even increased across all groups throughout the DSS cycle. This observation can be explained by the growth curve of Sprague-Dawley rats, which are still growing at eight weeks of age; therefore, disease manifestation may be reflected by stagnation or slower weight gain rate instead of weight loss. Similar findings have been previously reported (92,93). In the moderate inflammation model, both formulations consistently improved all disease-related endpoints. Histological analysis revealed preservation of the luminal epithelial lining, retention of goblet cells, and reduced crypt disruption in the LMP-treated groups. Additionally, mild inflammatory cell infiltration could explain the minor increase in tissue concentrations of TNF-α in the untreated group. In the severe inflammation model, the therapeutic benefit of the LMP-based formulations became even more evident at the macroscopic and biochemical levels, with prevention of colon shortening and marked reductions in DAI, fecal lactoferrin levels and tissue concentrations TNF-α.

TNF-α is a central mediator of intestinal inflammation and one of the most widely used molecular markers in the DSS model of colitis. DSS-induced epithelial injury leads to activation of innate immune pathways and subsequent production of pro-inflammatory cytokines, among which TNF-α plays a key role in amplifying mucosal damage, leukocyte recruitment, and barrier dysfunction (94,95). Numerous studies have demonstrated that TNF-α expression correlates with disease severity in DSS colitis models (96,97). Consistent with these reports, in our moderate colitis model, tissue concentrations of TNF-α remained comparable across all groups, indicating that inflammation at this stage is largely driven by epithelial injury and innate immune activation, rather than sustained TNF-α overproduction. In contrast, in the severe colitis model, TNF-α was a dominant inflammatory mediator, and both InflixiCaps and InflixiGel treatments markedly reduced its tissue levels toward those observed in naïve animals. This severity-dependent modulation of TNF-α aligns closely with previously described DSS models and supports the biological relevance of our system, demonstrating that therapeutic TNF-α neutralization becomes particularly effective under conditions of advanced mucosal damage and heightened inflammatory burden.

Despite macroscopic improvements, conventional histological scoring, primarily based on epithelial damage and immune cell infiltration, did not reveal significant differences between untreated and treated groups in the severe colitis model. The apparent mismatch between macroscopic improvement and histological scores under severe inflammation has been previously reported in DSS-induced colitis models (98–100) and likely reflects the inability of short-term anti-inflammatory treatments to fully reverse epithelial damage once severe injury has occurred. Chronic mucosal injury in UC has been reported to interrupt or even destroy the normal regeneration and repair processes of damaged intestinal epithelial cells (101,102), and the 6% DSS model induces particularly aggressive mucosal injury. Under these conditions, even the commercially available subcutaneous injection used as positive control was unable to reduce DAI, prevent colon shortening, or restore colonic epithelial integrity (Fig.S18, S19). Notably, both InflixiGel and InflixiCaps outperformed the systemic positive control, highlighting the advantage of localized IFX delivery in preserving mucosal barrier function under conditions of severe inflammation.

Importantly, the qualitative histological features preserved by InflixiGel and InflixiCaps (presence of continuous epithelial lining, retention of goblet cells, and reduction in crypt destruction) are mechanistically consistent with the known mucosal actions of IFX. In a seminal ex vivo and in vitro study, IFX was shown to directly promote mucosal healing (103). The preservation of goblet cells and the luminal epithelial layer observed in our LMP-treated animals, therefore, represents a biologically meaningful therapeutic outcome, as these structures are essential for preventing bleeding, bacterial translocation, and further inflammatory amplification in UC (101). In this context, the ability of InflixiGel and InflixiCaps to preserve goblet cells and epithelial continuity likely explains the marked reductions in fecal lactoferrin and disease activity observed in both moderate and severe models. These barrier preserving effects suggest that LMP-based colonic delivery not only suppresses inflammation but actively promotes mucosal healing, which is the strongest predictor of sustained remission and reduced hospitalization in UC patients. Previous studies have demonstrated that agents that promote hydration and restore colonic mucosal barrier function can improve UC outcomes by enhancing epithelial integrity and protecting the mucin layer (104). These observations underscore the importance of maintaining mucosal structure in mitigating inflammation and suggest that localized delivery strategies that preserve or protect the epithelial lining may offer therapeutic advantages.

## Conclusion

In this study, we demonstrate the previously underexplored potential of LMPs as versatile and effective platforms for targeted mAbs delivery to the inflamed colon. By doping MLO-based LMPs with SE, we successfully enlarged the aqueous channel diameter to enable efficient IFX release while simultaneously modulating gel rheology. This dual functionality proved critical for enabling SSE 3D printing of the oral dosage form InflixiPrint, and for facilitating the practical administration of the rectal formulation InflixiGel.

InflixiPrint exhibited gastro-resistance, prolonged matrix integrity and delayed, controlled release in the colon. In parallel, IFX release from InflixiGel was triggered by inflammatory conditions, a feature enabled by SE and highly relevant for targeting active disease sites.

Notably, LMP-based formulations successfully reached the colon following oral administration and exhibited strong mucoadhesion after rectal delivery. These properties translated into therapeutic benefits in a DSS-induced colitis model, with significant reductions in disease activity, inflammatory biomarkers, and histological damage. Overall, this study provides a new biocompatible and adaptable LMP-based delivery platform for colon-targeted drug delivery. By minimizing systemic exposure, while preserving therapeutic efficacy, this approach represents a promising advancement in the treatment of UC using biotherapeutics and provides a strong foundation for future translational development.

## Ethics statement

The work presented herein complies with all appropriate ethical regulations including animal welfare by the Cantonal Veterinary Offices of Berne, and the PREPARE and ARRIVE Guidelines.

## Study design

The goal of this study was to design a drug delivery system for UC. We hypothesize that the inclusion of a swelling agent can ameliorate the rheological properties and the delivery features of MLO-based mesophases. Thus, the developed formulations should rapidly adhere to the inflamed intestinal mucosa and release their cargo. Firstly, we characterized the formulations in vitro; subsequently, we examined whether mAb delivery via LMPs would exert efficacy in a DSS-induced colitis model in rats in compliance with institutional and federal regulations governing animal care and use, following the ARRIVE guidelines. Statistical details (sample size, outliers handling, randomization and blinding procedures, and strategies for comparisons among groups) for each in vivo experiment are provided in the figures’ captions, and a detailed description is reported in the Supplementary Information (SI).

## Methods

### Preparation of LMPs

The LMP composed of S80 and vitamin E (VitE) was prepared as previously described (33) and the detailed methodology is provided in the SI. The LMP composed of MLO and SE was prepared by melting and mixing the lipid with different percentages of SE (Table S2) in a magnetic stirrer for 1 h (500 rpm, 40°C). The MLO/SE lipid mixture (50% w/w) was hydrated with PBS or IFX solution (50% w/w) using the dual-syringe method as previously described (74), with detailed methodology provided in the SI. MLO_100_SE_0_ was produced by mixing MLO (60% w/w) and PBS or IFX solution (40% w/w). The preparation of IFX solutions and their quantification by HPLC and ELISA are described in the SI.

### Semi-solid extrusion 3D printing

The design of the 3D-printed tablets was created in AutoCAD 2024.1.2 (Autodesk, USA). The shell and core of the design were exported as .stl files, which were arranged in Slic3r, and the model was sliced to be printed with two LMPs. A Bio-X bioprinter (Cellink, Sweden) with a pneumatic thermoplastic printhead was used for 3D printing the tablets. To ensure optimal results, approximately 3 g of each formulation was placed into a pneumatic syringe and centrifuged at 4’430 x g for 15 min to remove air bubbles.

The optimal printing parameters were determined using a full factorial DoE generated by Minitab^®^ 18.1 software, with detailed methodology provided in the SI. The printing parameters for the S80/VitE LMP were pressure of 65 kPa, print speed of 3 mm/s, extruder temperature of 37°C, and nozzle diameter of 27G. The printing parameters for the MLO/SE LMP were pressure of 110 kPa, print speed of 3 mm/s, extruder temperature of 35°C, and nozzle diameter of 22G. For printing the core of the rat dosage form, a pressure of 160 kPa and a nozzle diameter of 20G were used. The tablets were stored at 2-8°C until use. The core was also printed separately to determine the core weight and, therefore, the correct dose. The weight of the full printlet was also determined.

### Rheology

A stress-controlled rheometer (Modular Compact Rheometer MCR 72, Anton Paar, Graz, Austria) was used in cone-plate geometry (0.993° angle and 49.942 mm diameter) to investigate the rheological properties of the LMPs. Viscosity-shear rate curves were measured at 25°C, and amplitude strain sweep measurements were performed at frequency of 10 Hz and strain percentages between 0.01 and 100% to determine both the LVR and the flow point.

### Small Angle X-Ray Scattering (SAXS)

SAXS measurements were performed on a Bruker AXS Micro using a microfocused X-ray source (λ Cu Kα = 1.5418 Å) operating at voltage of 50 kV and filament current of 1000 μA. The diffracted X-ray signals were collected using a 2D Pilatus 100 K detector. The scattering vector was calibrated using silver behenate. Data were collected and azimuthally averaged using the Saxsgui software to yield 1D intensity vs. scattering vector Q, with a Q range from 0.001 to 0.5 Å–1. For all measurements, the samples were placed inside a stainless-steel cell between two thin, replaceable mica sheets and sealed with an O-ring, with a sample volume of 10 μL and a thickness of ∼1 mm. Measurements were performed at 37°C, and the scattered intensity was collected over 30 min.

### Production of capsules

Gelatine capsules (size 4 for human dosing and size 9 for rat dosing) were filled with MLO_70_SE_30_ LMP and coated by dipping the capsules into Eudragit^®^ FS30D for 10 seconds and drying at room temperature for 10 minutes (five cycles). Capsules were filled with mannitol and IFX solution (10 µL, 80 mg/mL) and coated with Eudragit^®^ FS30D following the same procedure described above. The volume of the IFX solution was adjusted according to the weight of the rats to achieve an IFX dose of 2 mg/kg body weight.

### Disintegration studies of the oral dosage forms

Disintegration was assessed using an automated system (Agilent G7962A) in 900 mL of medium at 37°C. InflixiPrints (n=9) were incubated in HCl 0.1M for 1 h and then transferred to PBS, FaSSIF or FeSSIF (n=3). Disintegration was visually assessed at 15, 45, 75, and 105 min.

### Dissolution studies of the oral dosage forms

Dissolution experiments were performed using a basket apparatus (ERWEKA DT 126 light, Apparatus I). InflixiPrints were tested individually in 800 mL of medium at 37° C and 150 rpm, with incubation in HCl 0.1 M for 2 h followed by 6 h in PBS, FaSSIF, or FeSSIF (n=3). Samples were collected at the end of each phase, concentrated using 50’000 MWCO Amicon^®^ Ultra centrifugal filters, and IFX concentrations were determined by HPLC and ELISA. Using the same experimental setup, the dissolution profile of InflixiPrint was compared with that of coated capsules in HCl (2 h), FaSSIF (6 h), and FaSSCoF (1 h) (n=3).

### Lipolysis assay of the 3D-printed oral dosage form

In vitro lipolysis assays were performed following previously described methods with minor modifications (105), and the detailed methodology is provided in the SI.

### Release experiments of the rectal dosage form

InflixiGel (100 mg, 1.5 mg IFX) was weighed inside metallic baskets, placed inside falcon tubes containing 5 mL of release media (PBS with or without lipase (5000 U/mL), FaSSCoF with or without lipase (5000 U/mL) or FeSSCoF) and incubated at 37°C and 100 rpm for 8 h (n=3). The release medium was collected at 1, 2, 4 and 6 h and replaced with fresh 5 mL of buffer. The aliquots were concentrated in 50000 MWCO Amicon^®^ Ultra centrifugal filter and IFX concentration was determined by ELISA.

### Ex vivo experiments

Oral formulations (InflixiCaps or PowderCaps, one capsule inside each colon sample) and rectal formulations (200 mg of InflixiGel or 200 µL of enema inside each colon sample) were added inside colons samples (5 cm) from DSS-treated rats, 0.5 mL of FaSSCoF enriched with lipase was added, and both extremities were closed with surgical stitches (n=3 per treatment). Colon samples were incubated in FaSSCoF enriched with lipase (5000 U/mL) for 6 h at 37°C and 100 rpm. The colons were opened longitudinally, rinsed with 1 mL PBS, and homogenized with 2 mL PBS using a Polytron^®^ PT 2500 E homogenizer (Kinematica, Switzerland). Homogenates were centrifuged at 5’000 x g for 10 min at 4°C, and IFX concentration in the supernatants was quantified by ELISA according to the manufacturer’s instructions (HUMB00005, AssayGenie).

### In vivo experiments

Animal experiments were conducted using eight-week-old Sprague–Dawley rats (Janvier Laboratories, France) in accordance with the institutional and federal regulations governing animal care and use. The study complied with the PREPARE and ARRIVE guidelines and was approved by the Cantonal Veterinary Office of Bern (Switzerland) (BE89/2024). Detailed animal housing conditions are provided in the SI. Rats were randomly allocated to the different treatment groups on the experimental day. Each group included equal numbers of male and female animals, with sex included as a blocking factor in the statistical analysis. Because disease severity was sex-dependent, with males developing more severe disease, sexes were analyzed separately in figure 5 (panels C, L and N). Statistical details for each in vivo experiment, including sample size rationale, randomization, blinding, and exclusion criteria, are provided in the figure captions and described in detail in the SI.

### Progression of InflixiCaps and PowderCaps through the GI tract after oral gavage

GI transit following oral gavage was evaluated in healthy rats for InflixiCaps and PowderCaps loaded with barium sulfate (3 mg/capsule), incorporated either into the LMP or mannitol matrix, respectively. Capsule progression along the GI tract was assessed immediately after administration (t=0) and at 4 h and 6 h post-gavage. In preliminary studies, rats were administered capsules without an additional liquid contrast agent. The capsules were clearly observed but it was difficult to determine their location, as the GI tract was not highlighted. Therefore, 1 mL of a liquid contrast agent (Ultravist^®^ 300, Schering AG, Germany; 1 mL contains 0.623 g iopromide, 300 mg I/ mL) was given orally to rats by gavage under isoflurane anesthesia at different time points and administered together with the oral dosage forms. Oral gavage was performed using a rat capsule dosing tube (FTPU-C9-85, Instech).

Imaging: CT imaging was carried out using the SmART+ (Small Animal Radiotherapy System), Precision X-Ray, Inc. (Madison, CT, USA) equipped with a 0.4 mm focal spot X-ray source and calibrated to a source-to-isocenter distance of 30.7 cm and a source-to-detector distance of 64.5 cm. Animals were anesthetized with isoflurane (5% in 100% O_₂_ at 1 L/min) delivered via a nose cone and positioned in a stereotactic frame. For each scan, the SmART+ System performed a 360° rotation around the stationary animal, with the X-ray source and detector rotating together to acquire projections from multiple angles. These projections were reconstructed into a 3D cone-beam CT (CBCT) dataset, providing high-resolution anatomical information for accurate localization of the capsule within the GI tract.

### Colonic adhesion of InflixiGel after rectal administration

Healthy animals were fed with alfalfa-free diet 72 h before the experiment to suppress the autofluorescence in the abdominal region and reduce background signals. Immediately before acquiring the first imaging, the anogenital fur was shaved off to allow accurate dosing and imaging. InflixiGel and enema were loaded with DiD dye (0.02 mg/100 mg of formulation), and 150 mg of the formulation (n=6) was administered rectally to healthy animals under inhalation anesthesia (5 % isoflurane in 100 % O_2_ at 1 L/min). Rectal administration was performed by using sterile, disposable, flexible PTFE cannulas (Sigma CAD-9920) attached to a syringe containing the formulation. Animals were euthanized with a pentobarbital injection (150 mg/Kg intraperitoneal) at 8 h (n=3) and 24 h (n=3) post-rectal administration. Post-mortem, the colons were harvested and freshly imaged. Backgrounds (untreated tissue samples) were measured at each time point (n=3). The intensity of the fluorescent signal was measured using an IVIS SpectrumCT In Vivo Imaging System (PerkinElmer, MA, US) equipped with a 650 nm emission filter. The focus was kept stable using a subject height of 1.5 cm, and the temperature of the chamber was set at 37°C. Analysis of images (set with a fixed counts scale from 1’000 to 40’000 and acquired with a 0.2 s exposure time) were computed by first defining the region of interest (ROI) which was kept consistent across images and then the sum of all counts for all pixels inside the ROI (Total Fluorescence Counts- TFC; photons/second) recorded. Colonic adhesion profiles were obtained by plotting the TFC as a function of time (h).

### Blood and tissue pharmacokinetics

Blood: IFX concentration in the blood after administration of the formulations was evaluated in DSS-treated rats by giving free access to drinking water supplemented with 5.5% (w/v) DSS for 5 days, after which plain water was provided. Animals were randomly allocated to different treatment groups (n=6 rats/group), and on the 5^th^ day of DSS each animal received one of the following treatments: InflixiCaps, PowderCaps, InflixiGel, enema, subcutaneous injection. Dose for rectal formulations was 5 mg/kg, and for subcutaneous injection and oral formulations was 2 mg/kg. Blood (110 µL) was collected in BD Microtainer^®^ serum tubes from the lateral tail vein at 0.5, 4, 8, 24 h, 2, 4, 7, 15, and 22 days after treatment administration and allowed to clot by leaving it undisturbed at room temperature for 15 min. Samples were centrifuged at 4°C and 1’500 x g for 10 minutes. The serum was collected and stored in cryotubes at -20°C until drug quantification by ELISA according to manufacturer’s instructions (HUMB00005, AssayGenie). AUC 0-22d was calculated using a noncompartmental analysis with intravascular administration as a model. After the last sampling point, animals were euthanized with CO_2,_ and the colon was collected for tissue IFX quantification.

Tissue: Colonic retention of IFX following oral or rectal administration was evaluated in healthy and DSS-treated rats. Colitis was induced by giving free access to drinking water supplemented with 5.5% (w/v) DSS for 5 days. On the 5^th^ day of DSS, colitic rats were randomly assigned to one of the following treatment groups: InflixiCaps, PowderCaps, InflixiGel, enema, or subcutaneous IFX injection. Animals were euthanized at predefined time points using pentobarbital (150 mg/kg, intraperitoneal), and colonic tissues and luminal contents were collected for IFX quantification. Rats receiving oral formulations (InflixiCaps or PowderCaps) were euthanized 24 h after gavage (n=4 rats/group). Rats receiving rectal formulations (InflixiGel or enema) were euthanized at 12 and 24 h after administration (n=4/group). Animals receiving subcutaneous IFX were euthanized at 7 and 14 days post-injection (n=4/group). Healthy rats were randomly assigned to receive InflixiCaps, PowderCaps, InflixiGel, or an enema. Animals receiving oral formulations were euthanized at 24 and 48 h after gavage (n=3/group), whereas animals receiving rectal formulations were euthanized at 12 and 24 h post-administration (n=3/group).

Following euthanasia, stool samples were collected, suspended in PBS containing protease inhibitors (cOmplete™, Roche Diagnostics, Germany), vortexed, and centrifuged at 5’000 x g for 10 min at 4°C. Supernatants were stored at −20°C until analysis. Colons were rinsed with PBS, cut into small fragments, and homogenized in PBS containing protease inhibitors using a Polytron^®^ PT 2500 E homogenizer (Kinematica, Switzerland). Homogenates were centrifuged at 5’000 x g for 10 min at 4°C, and the supernatants were concentrated using 50’000 MWCO Amicon^®^ Ultra centrifugal filters before storage at −20°C. IFX concentrations in stool and tissue extracts were quantified by ELISA according to the manufacturer’s instructions (HUMB00005, AssayGenie).

### Therapeutic efficacy evaluation following a prophylactic approach

Prophylactic therapeutic efficacy of oral and rectal formulations was evaluated in a DSS-induced colitis model in rats. Moderate and severe colitis were induced by providing free access to drinking water supplemented with 4% and 6% (w/v) DSS for seven days, respectively. Animals were randomly allocated to different treatment groups (n=6 rats/group), and on the first day of DSS supplementation each animal received one of the following treatments: subcutaneous injection (single administration, 2 mg IFX/kg), InflixiCaps or PowderCaps (daily administration for seven consecutive days, 2 mg IFX/kg), InflixiGel or enema (daily administration for seven consecutive days, 5 mg IFX/kg). Following each administration, animals were monitored for 1 h. In the rectally treated groups, if formulation excretion was observed within this period, a second dose was administered to ensure effective colonic exposure. Six animals received DSS supplementation and no treatment and were used as a control. DAI was assessed daily and calculated based on inflammation associated parameters: body weight loss (percentage of weight loss relative to the initial body weight, where 0, <3%; 1, 3-<10%; 2, 10-<15%; 3, ≥15%), stool consistency (0, normal; 1, sticky and/or moist; 2 mild-to-moderate diarrhea; 3 severe diarrhea), and hematochezia (0, no blood; 1, traces of blood in the stool; 2, bloody stool and/or blood around the anus; 3, severe bleeding). On the 8^th^ day, rats were euthanized with intraperitoneal pentobarbital injection (150 mg/kg), colon length was determined, and colon samples were collected for histochemical analysis and TNF-α quantification. Additionally, stool samples were collected and lactoferrin concentration was quantified by ELISA following manufacturer’s instructions (RTFI00940, AssayGenie).

#### TNF-α quantification

The second most distal 1 cm of the colon was collected for TNF-α quantification. Colon pieces were lysed with cell lysis buffer (100 µL/ 5 mg tissue, Bio-Rad) using a GentleMACS device from Miltenyi Biotec (Miltenyi Biotec, Bergisch Gladbach, Germany), and the concentration of TNF-α in the lysates was determined by ELISA following manufacturer’s instructions (AEFI01710, AssayGenie).

#### Histology

To assess the microscopic extent of colitis, the most distal part of the colon (1 cm) was fixed in 4% formalin and subsequently embedded in paraffin. Sections of 5 µm were cut using a rotary microtome (Zeiss, Oberkochen, Germany) and deparaffinized with Histoclear (Chemie Brungschwig, Basel, Switzerland). Sections were rehydrated using a descending series of ethanol (100 to 70%) and stained with hematoxylin for 10 min followed by a 2 s differentiating step with 1% HCl and further stained for 10 sec with eosin. Sections were dehydrated with an ascending series of ethanol (70 to 100%) and mounted with Pertex (HistoLab, Askim, Sweden). Histological scoring was performed by three independent investigators blinded to the type of treatment, and epithelial damage and inflammatory cell infiltration were assessed (106). Detailed description of the histology scoring system is provided in the SI.

#### Histological quantification of colonic morphology

Quantitative histological analysis was performed on the distal colon to assess epithelial integrity (107,108). The most distal part of the colon (1 cm) was analyzed for each animal using a systematic uniform random sampling approach. For each sample, three non-overlapping fields were selected, each consisting of a stitched (patchwork) image generated from 10 high-magnification micrographs arranged in three rows. The area of preserved luminal epithelial lining was measured within each stitched field, and the measurements from the three fields were pooled. Data were expressed as the percentage of preserved luminal epithelial lining in the 1 cm colon sample and then summarized per treatment group and sex for statistical analysis. This approach ensured unbiased sampling and robust quantification of histological features across the distal colon.

## Acknowledgments

Dr. Laura Baraldi (Laboratory of Food & Soft Materials, Institute of Food, Nutrition and Health, IFNH; Department for Health Sciences and Technology, D-HEST, ETH Zurich Switzerland) is kindly acknowledged for her support during the SAXS experiments. Andreas Niederquell (Institute of Pharma Technology, University of Applied Sciences and Arts Northwestern Switzerland) is kindly acknowledged for his support during the lipolysis experiments. Dr. Aurelie Quintin (Medova Lab, Department of Biomedical Research, University of Bern, Switzerland)) and Dr. Julien Ott (Inselspital, University Hospital Bern, Switzerland) are kindly acknowledged for their support during the computed tomography imaging. Doris Pöhlmann (Gastroscience Scharl Lab, Department of Gastroenterology and Hepatology, University Hospital Zurich/University of Zurich, Switzerland) and Vanessa Dobernig (Rogler Lab, Department of Gastroenterology and Hepatology, University Hospital Zurich/University of Zurich, Switzerland) are kindly acknowledged for their support during the histology experiments. Dr. Peter Eguia, Dr. Timokleia Kousi, Sandro Grepper and Eva Zwygart (Experimental Animal Center, University of Bern, Switzerland) are kindly acknowledged for their support during the animal experiments. The authors gratefully acknowledge Swiss National Science Foundation for the financial support (Grant Nr. 10.003.897).

## Authorship contribution statement

**Rafaela Gazzi:** Writing – original draft, Writing – review & editing, Visualization, Validation, Methodology, Investigation, Formal analysis, Data curation, Conceptualization. **Tiziana Cremona:** Writing – original draft, Writing – review & editing, Validation, Methodology, Investigation, Formal analysis, Data curation, Conceptualization**. Reto Czekalla:** Investigation, Formal analysis. **Emily Campbell:** Writing – review & editing, Investigation, Formal analysis. **Marlene Schwarzfischer:** Writing – review & editing, Investigation, Formal analysis. **Gerhard Rogler:** Writing – review & editing, Resources. **Michael Scharl** Writing – review & editing, Resources. **Alessandra Bergadano** Writing – review & editing. **Raffaele Mezzenga** Writing – review & editing, Formal analysis (SAXS), Resources. **Martin Kuentz** Writing – review & editing, Resources. **Paola Luciani** Writing – review & editing, Resources, Supervision, Funding Acquisition. **Simone Aleandri** Writing – original draft, Writing – review & editing, Methodology, Investigation, Formal analysis, Data curation, Conceptualization, Supervision, Funding Acquisition, Project Administration.

