## Supplementary Information for "Shifting Perspectives on Biotherapeutic Treatment of Ulcerative Colitis using Lipid Mesophases: Formulation Design and Preclinical Validation"

**Affiliations:**

### Supplementary Materials and Methods

#### Materials:

S80 (with 75% polyenylphosphatidylcholine) and Lipoid E PC S (phosphatidylcholine from egg yolk) were obtained from Lipoid GmbH (Ludwigshafen, Germany). Our source of monolinolein was Dimodan U/J, which was kindly donated by Danisco (Copenhagen, Denmark). Dimodan U/J is an industrial food-grade quality analogue of MLO composed of a mixture of monoglyceride derived from oleic fatty acid with a minimum monoglyceride content of 90%. Sucrose stearate was purchased from abcr (Karlsruhe, Germany). Vitamin E (Ph. Eur. Quality), lipase from Candida rugosa, Trizma® maleate, calcium chloride dihydrate (≥99%), pancreatin (from porcine pancreas, 8xUSP specifications) and maleic acid were purchased from Sigma-Aldrich Chemie GmbH (Buchs, Switzerland). Phosphate buffer saline was obtained from Capricorn Scientific GmbH (Ebsdorfergrund, Germany). Disodium phosphate, embedding cassettes and sponges for histology and formaldehyde solution (37%) were obtained from Carl Roth (Karlsruhe, Germany). Sodium dihydrogen phosphate dihydrate was obtained from Hänseler AG (Herisau, Switzerland). Sodium taurodeoxycholate was obtained from Prodotti Chimici e Alimentari S.p.A. (Basaluzzo, Italy). Barium sulphate (Puratronic, 99.998%) was purchased from Thermo Fisher Scientific (Switzerland). Hydrochloric acid 37% pure (HCl) was supplied by Dr. Grogg Chemie (Switzerland). Eudragit® FS30D was kindly donated by Evonik (Rheinfelden, Germany). Simulated digestive fluids were purchased from Biorelevant Ltd. (London, UK). Infliximab (IFX, Remicade) was purchased from the Inselspital (Bern, Switzerland). Infliximab (HUMB00005), rat lactoferrin (RTFI00940) and rat TNF-α (AEFI01710) ELISA kits were obtained from AssayGenie (Dublin, Ireland). 1,1′-dioctadecyl-3,3,3′,3′-tetramethylindodicarbocyanine, 4chlorobenzenesulfonate salt (DiD) was from Invitrogen (Carlsbad, USA). Gelatine capsules size 4 were from purchased from Interdelta (Switzerland), and gelatine capsules size 9 were purchased from Torpac (Heerlen, Netherlands). Dextran sodium sulphate (DSS, 40kDa) was obtained from TdB Labs (Uppsala, Sweden). Cell lysis kit was purchased from Bio-Rad (Switzerland).

#### Preparation of LMPs:

*S80/VitE LMP*

S80 (80%) and vitamin E (12%) were co-dissolved in methanol into a glass vial, and the organic solvent was removed under reduced pressure (0.22 mbar for 24 h). Water (20%) was incorporated into the dried lipid, and the mixture was manually mixed with a spatula and centrifuged for 10 min at 5’000 x g, which was repeated six times to obtain a uniform and homogeneous gel. The LMP was allowed to equilibrate for 48 h in the dark at room temperature.

*Preparation of MLO/SE LMP*

Monolinolein (MLO) and sucrose stearate (SE) mixture (50% w/w) was transferred to a syringe (Fisher Scientific, Pittsburgh, US), and the aqueous solution (PBS or IFX solution, 50%) was added to a second syringe. Both syringes were attached to a syringe connector (Combifix® Adapter Luer-Lock female/female), and the LMP was hydrated by transferring the contents of one syringe to the other for three min. The final gel was equilibrated in the syringe for 10 min.

**Infliximab solution preparation and quantification**

Remicade vials were resuspended in ultrapure water to a concentration of 30 mg/mL and used to produce the rectal gels. For comparison in vivo, the solution was diluted to 15 mg/mL and administered as an enema. The 30 mg/mL IFX solution was concentrated using a 50'000 MWCO Amicon® Ultra centrifugal filter to 100 mg/mL to print the core of the human printlet, and to 180 mg/mL to print the core of the rat printlet. The concentration of IFX in preliminary release studies in PBS and simulated intestinal fluids was determined by HPLC as described below. The concentration of IFX in confirmatory release studies in simulated intestinal fluids and in vivo experiments was determined by ELISA (HUMB00005, AssayGenie) following manufacturer's instructions.

**HPLC method for IFX quantification**

IFX concentrations were determined by reversed-phase high-performance liquid chromatography (RP-HPLC) using a Biozen dSEC 2 (00H-4788-K0, Phenomenex, USA). The HPLC method was adopted from the European Pharmacopoeia, with a flow rate of 1 mm/s, column temperature of 25°C, detection wavelength of 214 nm, injection volume of 10 µL, and run time of 20 min. The sampler was kept at 10°C. The mobile phase was composed of 6.5 mM $NaH_2PO_4$, 13.5 mM $Na_2HPO_4$, and 150 mM NaCl.

**Circular dichroism**

To determine if the concentration of IFX affects its structure, circular dichroism measurements were performed on IFX solutions with concentrations of 30, 100, 180 mg/mL. A J-75 detector coupled with a PS150J power unit (JASCO, Germany) was used. The resulting spectra were then compared to IFX spectra from the literature to determine if the high concentration has an influence on the structure. The solutions were diluted to 0.05 mg/mL for successful measurements. Cuvettes with a 1 cm path length were used for measurements between 205 and 250 nm at room temperature.

**Design of Experiments:**

The optimal printing parameters were determined using a full factorial DoE generated by Minitab® 18.1 software. Extruder temperature (three levels: 30, 35, and 40°C), speed (three levels: 3, 5, and 7 mm/s), and pressure (four levels: 80, 110, 150, and 200 kPa) were selected as factors. In total, 36 runs were performed for each of the six LMPs composed of different percentages of MLO and SE, and a printability score (from 0 to 5, where 0 was not possible to print and 5 the optimal printlet shape and consistency) was used as response. A scoring guide for printability is shown in Figure S1. From the printability scores, a model was calculated to determine the influence of factors on the results, and which factors have statistically significant effects on the response ($p \leq 0.10$).

**Lipolysis assay of the 3D-printed oral dosage form**

A micellar solution simulating fasted-state intestinal conditions was prepared by dissolving phosphatidylcholine (1.25 mM) and sodium taurodeoxycholate (5 mM) in a digestion buffer (50 mM Trizma® maleate, 150 mM NaCl, 5 mM $CaCl_2 \cdot 2H_2O$, pH 7.5). The mixture was stirred for 12 h at 5°C (450 rpm). A pancreatin extract was freshly prepared by dispersing porcine pancreatin powder (1 g/5 mL digestion buffer), stirring for 15 min at 5°C, and centrifuging (15 min, 1600 x g, 5°C). The supernatant pH was adjusted to 7.5 and kept on ice until further use. The lipolysis experiment was conducted by adding one 3D-printed tablet to 60 mL of the micellar medium in a thermostated vessel (37 ± 0.5°C) under stirring (700 rpm). After 10 min of equilibration, digestion was initiated by adding pancreatin extract (final nominal lipase activity: 1000 tributyrin units/mL). Free FA released during digestion were titrated automatically with 1 M NaOH to maintain a pH of 7.5 using a pH-stat apparatus (842 Titrando and 800 Dosino, Metrohm AG, Switzerland). Lipolysis proceeded for 270 min. To correct for unionized free FA, back-titration was performed by rapidly increasing the pH to 9 with 1 M NaOH at the end of digestion. A blank digestion (without formulation) was also performed, and its NaOH consumption was subtracted to obtain the corrected free FA value (FAtitr). The correction factor was calculated according to the formula:

Correction factor = (FAtitr (direct titration) + FAtitr (back titration)) / FAtitr (direct titration)

where FAtitr (direct titration) is the amount of FA titrated after the 270 min digestion period.

A correction factor was calculated for both the lipolysis of the 3D-printed tablet and pure digestion medium. The amount of free FA released from the pure digestion medium was then subtracted from the total FA measured in the presence of the tablet.

**In vivo experiments**

Rats were housed under specific pathogen-free conditions in compliance with the Federation of European Laboratory Animal Science Associations (FELASA) guidelines. Animals were kept in groups of up to three in autoclaved individually ventilated cages (Blue Line, 1500 $cm^2$, Tecniplast, Italy) containing aspen wood bedding (J. Rettenmeier & Söhne GmbH, Germany) and cotton nestlets that were used for nesting. Environmental enrichment was provided in the form of a black PVC tunnel (Plexx, Netherlands) and an aspen wood stick (LAB & VET Service GmbH, Austria). Autoclaved tap water and irradiated rodent chow (Mouse and Rat Maintenance 3432; Granovit, Switzerland) were provided ad libitum. Housing conditions were maintained under a 12:12 h light-dark cycle, at a room temperature of 22 ± 2°C and a relative humidity of 45-65%. Animals were allowed a one-week acclimatization period and were regularly handled by trained personnel for gentling and habituation to the experimental procedures.

**Histology scoring**

Histological scoring was performed by three independent investigators blinded to the type of treatment, and epithelial damage and inflammatory cell infiltration were assessed.
Epithelial damage: intact epithelium = 0; focal goblet cell depletion in limited areas and/or mild irregularity in crypt spacing = 1; moderate goblet cell loss across larger areas and/or pronounced irregular crypt spacing = 2; extensive loss of goblet cells and crypt structures = 3; near complete or complete loss of crypts over large areas = 4.
Infiltration: no infiltrate = 0; mild infiltration around crypt basis = 1; moderate infiltration extending into the submucosa = 2; extensive infiltration throughout the submucosa and oedema = 3; dense infiltration into both submucosa and muscularis externa, prominent oedema and tissue disruption = 4.
The total histology score represents the sum of the epithelium damage score and the infiltration score. Images were taken using a Zeiss Axio Imager.Z2 microscope (Zeiss), equipped with an AxioCam HRc (Zeiss, Jena, Germany) camera and ZEN imaging software (Zeiss, Germany).

**Statistical analysis performed for the in vivo studies:**

| Imaging studies (Figure 3; Panel E) | |
|---|---|
| Sample size (n) | The significance threshold was set at a value of $p<0.05$, and the desired power is 0.9. An effect size (d; difference between means/ standard deviation) set at 0.5 would already be clinically relevant. In such conditions, when comparing 2 groups, 6 samples/group are required. F test; ANOVA: repeated measurements, within-between interaction. A priori: Compute the required sample size. The test was performed within the G*power software |
| Randomization | On arrival from the company, animals were randomized using a computer-based random order generator and the treatments were randomly assigned to the animals within the cage. |
| Blinding | The experimenter was blinded regarding the administered formulations (performed by a second operator) while assessing the imaging study. |
| Statistical methods | The results are presented as mean ± SD. Multiple comparison between the groups was performed by a parametric test (repeated measures ANOVA). A post-hoc correction for multiple comparison was done using Tukey.<br><br>Only descriptive statistics were used during the evaluation of transit time (Figure 3, Panel B). The scope of this aim is not a comparison between groups. |

| PK studies (Figure 4; Panels G-L) | |
|---|---|
| Sample size (n) | The scope of this aim is not to draw a general conclusion about IFX systemic absorption (PK population) in rats, which will require a higher sample size to represent the population, but a comparison study for the drug's systemic absorption after different administration routes. The significance threshold was set at a value of $p<0.05$, and the desired power is 0.9. An effect size (d; difference between means/ standard deviation) set at 0.25 would already be clinically relevant. In such conditions, 6 samples/group are required for a parametric test. F test; ANOVA: repeated measurements, within-between interaction. A priori: Compute the required sample size. The test was performed within the G*power software. |
| Randomization | On arrival from the company, animals were randomized using a computer based random order generator and assigned to the different blocks. The treatments were randomly assigned to the animals within the block. |

| | |
|---|---|
| Blinding | The experimenter was blinded regarding the administered formulations (performed by a second operator) while assessing the blood sampling. |
| Statistical methods | The results are presented as mean ± SD. Comparison between the groups was performed by a parametric test (repeated measures two-way ANOVA). A post-hoc correction for multiple comparison was done using Tukey test. |

| PD studies (Figure 5; Panels A-F and K-Q) | |
|---|---|
| Study design | The study plan shows a comparative study for the evaluation of anti-inflammatory efficacy after rectal and oral administration of treatments. Due to the difficulties in handling the total sample size, the experiment was divided into small blocks and the experiment on each block was carried out in a different time (the execution day of the experiment and sex was be used as blocking factor). |
| Sample size (n) | The significance threshold was set at a value of $p<0.05$, and the desired power is 0.90. An effect size (d; difference between means/ standard deviation) set at 0.25 would already be clinically relevant. In such conditions, 6 samples/group are required. F test; ANOVA: repeated measurements, within-between interaction. A priori: Compute the required sample size. The test was performed within the G*power software. |
| Randomization | Randomization was used to create groups containing an equal number of animals administered with each treatment (1 animal/ treatment) and 1 untreated animal (used as control group). On arrival from the company, animals were randomized using a computer based random order generator and assigned to the 6 different blocks. The treatments were randomly assigned to the animals within the block. |
| Blinding | The experimenter was blinded regarding the administered formulations (performed by a second operator) while evaluating the read out. |
| Statistical methods | The results are presented as mean ± SD or mean ± SEM. Depending on the results, multiple comparisons between the groups were analyzed by:<br><br>- Repeated measures (DAI see Figure 5, Panels A, K and L): a linear mixed-effects model (LME) was used. The LME was selected to account for the repeated measurements taken from individual animals over time (random effect) and to handle the unbalanced nature of the dataset. Fixed effects included Group, Time, and their interaction.<br><br>- One-way ANOVA for the comparison of all the other readouts and a post-hoc correction for multiple comparison was done using Tukey test. |

### Supplementary Tables

**Supplementary Table 1.** Printability scores for DoE runs for LMPs with lipid mixture containing 0-40% sucrose stearate.

| Factors | | | Printability score of tested LMPs | | | | | |
|---|---|---|---|---|---|---|---|---|
| Pressure (kPa) | Speed (mm/s) | Temperature (°C) | 0 % LMP | 20 % LMP | 25 % LMP | 30 % LMP | 35 % LMP | 40 % LMP |
| 150 | 7 | 40 | 0 | 1 | 1 | 1 | 1 | 1 |
| 150 | 5 | 40 | 0 | 1 | 1 | 1 | 1 | 1 |
| 200 | 5 | 40 | 0 | 1 | 1 | 1 | 1 | 1 |
| 200 | 5 | 35 | 0 | 2 | 1 | 1 | 1 | 1 |
| 110 | 5 | 30 | 0 | 1 | 1 | 4 | 1 | 1 |
| 80 | 5 | 35 | 0 | 1 | 1 | 2 | 4 | 2 |
| 110 | 5 | 40 | 0 | 1 | 1 | 2 | 1 | 1 |
| 200 | 7 | 30 | 0 | 1 | 1 | 1 | 1 | 1 |
| 150 | 3 | 30 | 0 | 1 | 3 | 1 | 1 | 1 |
| 150 | 7 | 30 | 0 | 1 | 1 | 1 | 1 | 1 |
| 110 | 7 | 35 | 0 | 1 | 1 | 2 | 2 | 1 |
| 80 | 5 | 40 | 0 | 1 | 1 | 1 | 1 | 3 |
| 200 | 7 | 40 | 0 | 2 | 1 | 1 | 1 | 1 |
| 80 | 3 | 35 | 0 | 1 | 1 | 2 | 4 | 1 |
| 200 | 3 | 30 | 1 | 1 | 1 | 1 | 1 | 1 |
| 110 | 3 | 30 | 0 | 1 | 1 | 3 | 1 | 1 |
| 110 | 3 | 40 | 0 | 1 | 1 | 4 | 1 | 1 |
| 200 | 5 | 30 | 0 | 1 | 1 | 1 | 1 | 1 |
| 80 | 7 | 35 | 0 | 1 | 1 | 1 | 2 | 2 |
| 200 | 7 | 35 | 0 | 2 | 1 | 1 | 1 | 1 |
| 80 | 5 | 30 | 0 | 1 | 1 | 1 | 4 | 1 |
| 200 | 3 | 40 | 1 | 1 | 1 | 1 | 1 | 1 |
| 80 | 3 | 40 | 0 | 1 | 1 | 1 | 3 | 1 |
| 110 | 5 | 35 | 0 | 1 | 1 | 3 | 1 | 1 |
| 80 | 3 | 30 | 0 | 1 | 1 | 1 | 5 | 1 |
| 110 | 7 | 40 | 0 | 1 | 1 | 1 | 2 | 1 |
| 110 | 3 | 35 | 0 | 1 | 1 | 5 | 1 | 1 |
| 110 | 7 | 30 | 0 | 1 | 1 | 3 | 1 | 1 |
| 150 | 3 | 35 | 0 | 1 | 2 | 1 | 1 | 1 |
| 80 | 7 | 40 | 0 | 1 | 1 | 1 | 1 | 4 |
| 150 | 5 | 35 | 0 | 1 | 2 | 1 | 1 | 1 |
| 150 | 5 | 30 | 0 | 1 | 2 | 1 | 1 | 1 |
| 80 | 7 | 30 | 0 | 1 | 1 | 1 | 4 | 2 |
| 200 | 3 | 35 | 1 | 1 | 1 | 1 | 1 | 1 |
| 150 | 7 | 35 | 0 | 1 | 1 | 1 | 1 | 1 |
| 150 | 3 | 40 | 0 | 1 | 1 | 1 | 1 | 1 |

**Supplementary Table 2.** Ratio of lipid (MLO) and sucrose stearate (SE) in the lipid mixture and the corresponding LMP nomenclature.

| LMP Nomenclature | Monolinolein (%) | SE (%) |
|---|---|---|
| $MLO_{100}SE_{0}$ | 100 | 0 |
| $MLO_{80}SE_{20}$ | 80 | 20 |
| $MLO_{75}SE_{25}$ | 75 | 25 |
| $MLO_{70}SE_{30}$ | 70 | 30 |
| $MLO_{65}SE_{35}$ | 65 | 35 |
| $MLO_{60}SE_{40}$ | 60 | 40 |

### Supplementary Figures

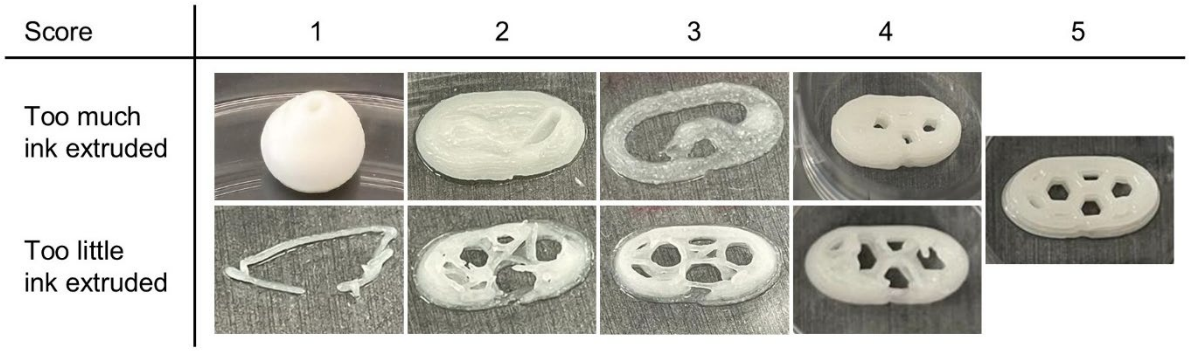


**Supplementary Figure 1.** Printability scoring guide for the DoE runs. Scores 1-4 correspond to printing conditions that produce excessive (top) or insufficient (bottom) ink extrusion. Score 5 indicates an optimal 3D-printed core. Score 0 is assigned when no ink is extruded from the nozzle (not shown).

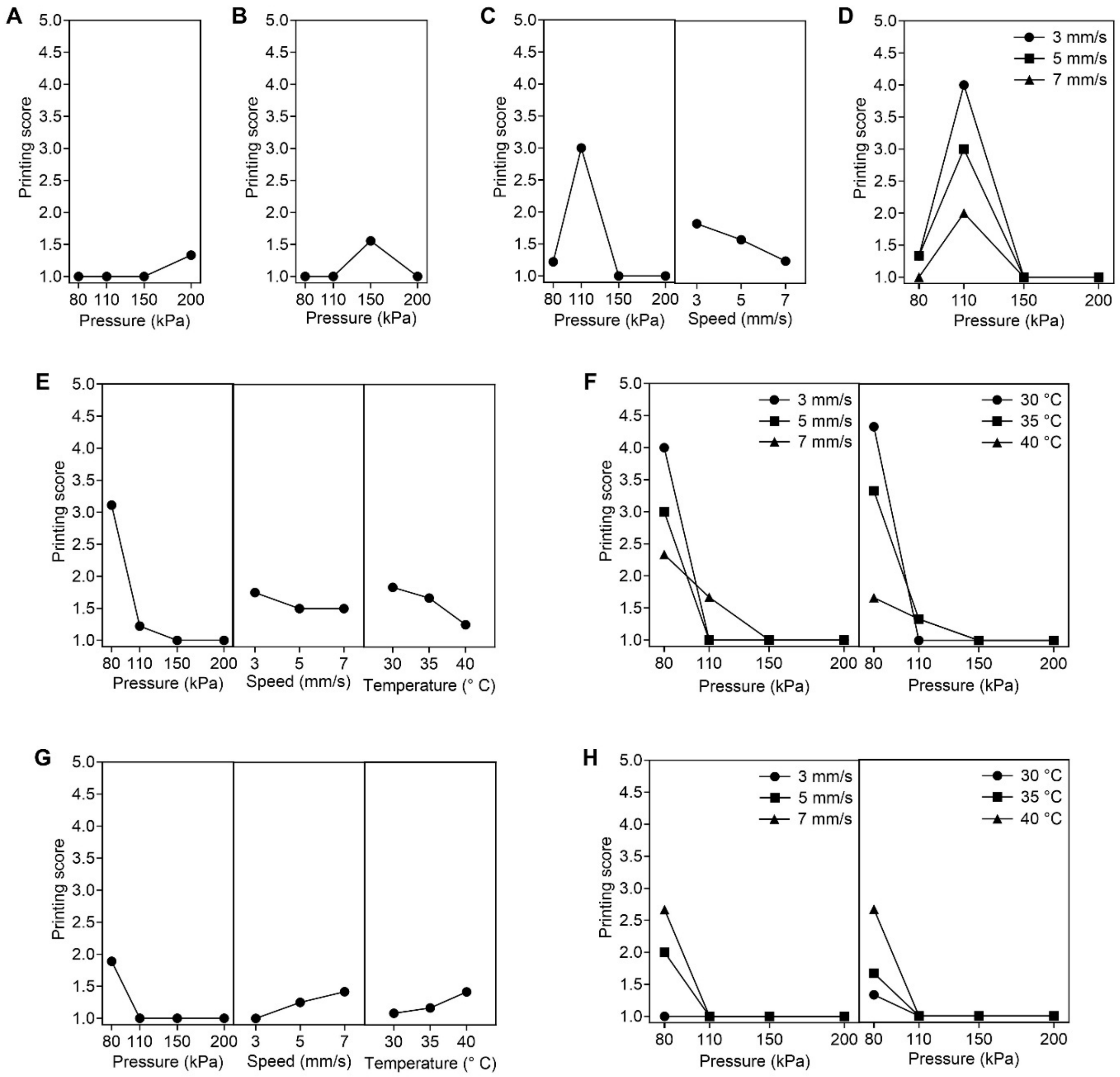


**Supplementary Figure 2.** Main effects and interaction plots for printability of LMPs with sucrose stearate 20-40%. Main effects plots for the printability of (A) $MLO_{80}SE_{20}$ and (B) $MLO_{75}SE_{25}$. (C) Main effects and (D) interaction plots (pressure × speed) for the printability of $MLO_{70}SE_{30}$. (E) Main effects and (F) interaction plots (left: pressure × speed, right: pressure × temperature) for printability of $MLO_{65}SE_{35}$. (G) Main effects and (H) interaction (left: pressure × speed, right: pressure × temperature) plots for printability of $MLO_{60}SE_{40}$.

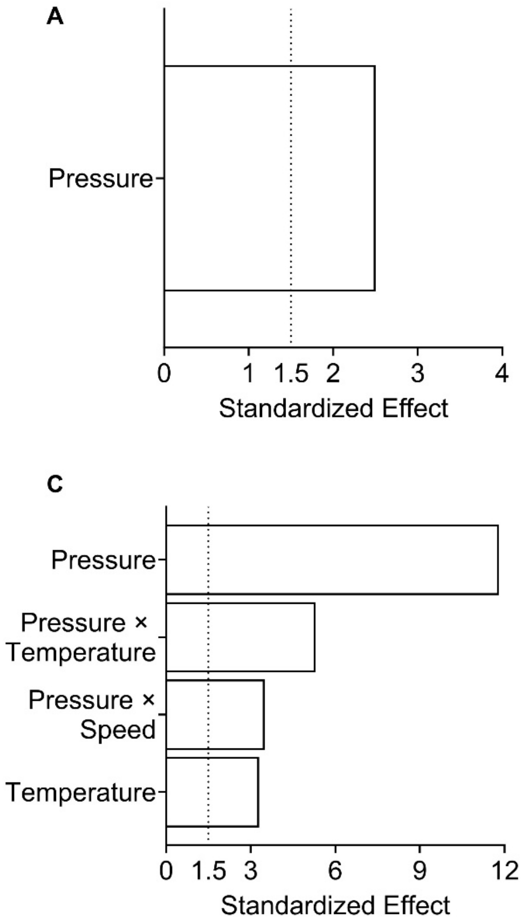


**Supplementary Figure 3.** Pareto charts of the standardized effects on the printability of LMPs with sucrose stearate at 20-40%. (A) $MLO_{80}SE_{20}$, (B) $MLO_{75}SE_{25}$, (C) $MLO_{65}SE_{35}$ and (D) $MLO_{60}SE_{40}$.

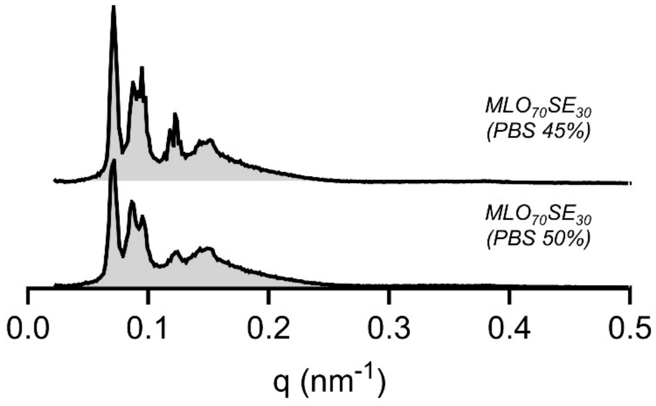


**Supplementary Figure 4.** SAXS spectra of $MLO_{70}SE_{30}$ hydrated with 45% or 50% PBS. Both spectra show Bragg peaks with positions in the ratio √2:√3:√4:√6:√8:√9, characteristic of an inverse bicontinuous cubic phase with Pn3m symmetry. Both LMPs present the first Bragg peak (√2) at q 0.07 $nm^{-1}$. The LMP is at maximum hydration as the first Bragg peak (√2) does not change its position while increasing the hydration percentage hydration. In contrast, $MLO_{100}SE_0$ reaches its maximum hydration with 40% of PBS and its first Bragg peak is at q > 0.15 $nm^{-1}$ (see Fig. 1, panel K upper spectra).

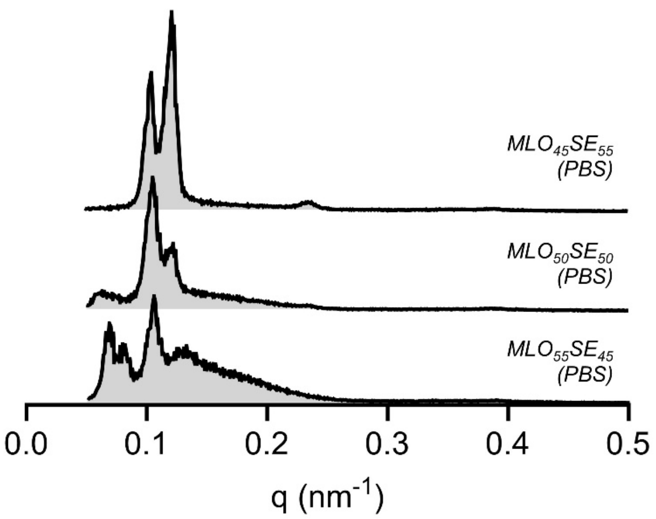


**Supplementary Figure 5.** SAXS spectra of LMPs composed of 50% lipid mixture of MLO and sucrose stearate (SE) and 50% PBS. Top spectra: lipid mixture composed of 45% MLO and 55% SE. Middle spectra: Lipid mixture composed of 50% MLO and 50% SE. Bottom spectra: Lipid mixture composed of 55% MLO and 45% SE. These percentages of SE lead to LMPs with unknown crystalline domains.

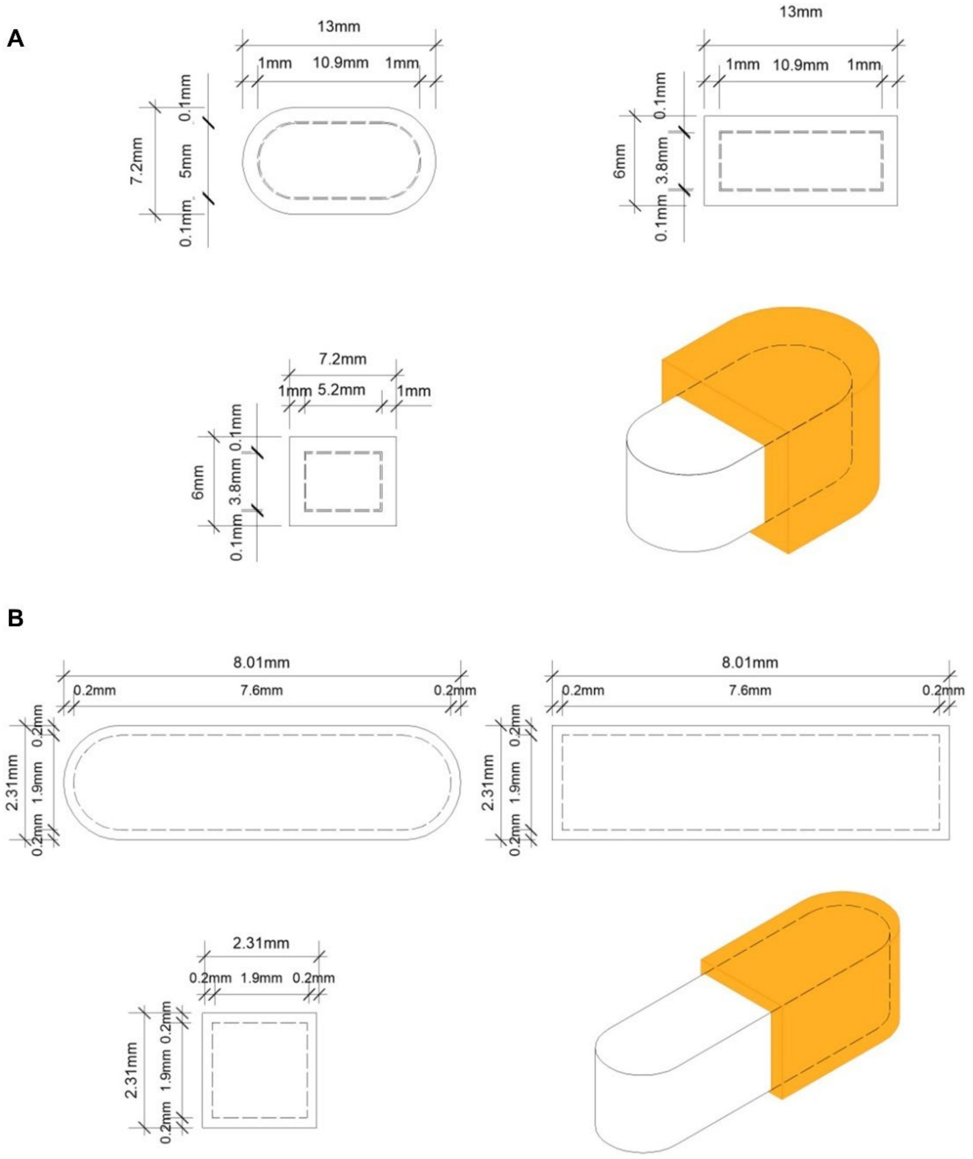


**Supplementary Figure 6.** Technical drawings of the designed 3D-printed oral dosage forms. (A) Tablet designed for human use. (B) Scaled-down tablet for in vivo experiments in rats.

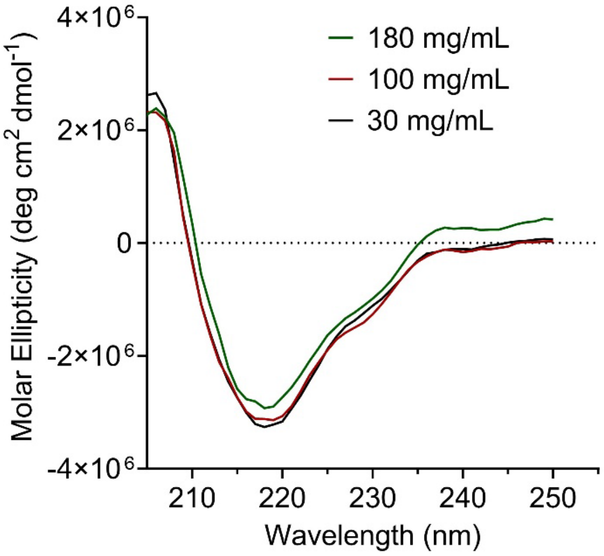


**Supplementary Figure 7.** Far-UV circular dichroism spectra of infliximab solutions at 30, 100, and 180 mg/mL. A strong negative band near 217 nm, indicative of the β-sheet secondary structure, was consistent across all concentrations.

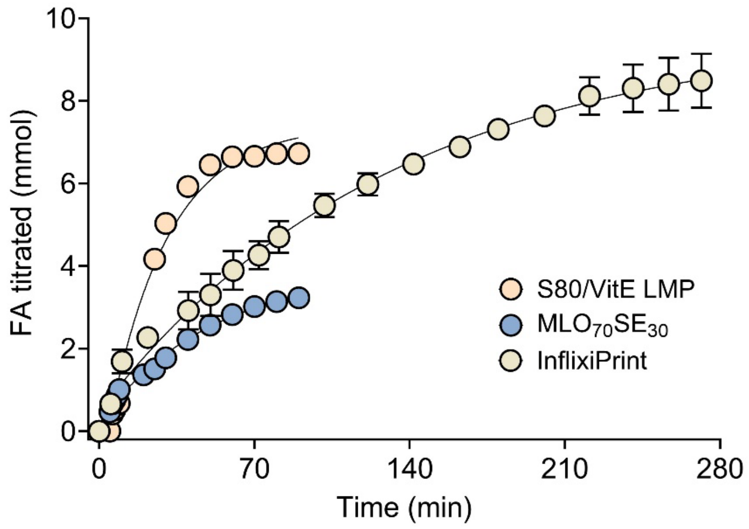


**Supplementary Figure 8.** Free fatty acids (FA) originated from the digestion of InflixiPrint (n=2) or its individual components (S80/VitE LMP in the shell or $MLO_{70}SE_{30}$ LMP in the core).

A B

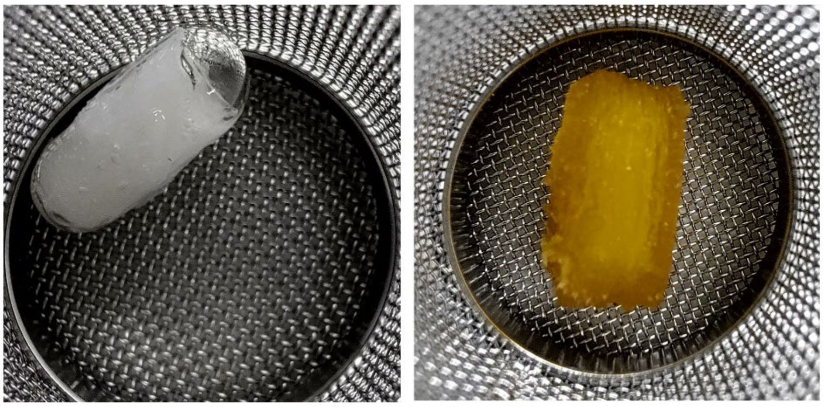

**Supplementary Figure 9.** Representative images of (A) InflixiCaps and (B) InflixiPrint after 2 h of incubation in HCl (0.1 M).

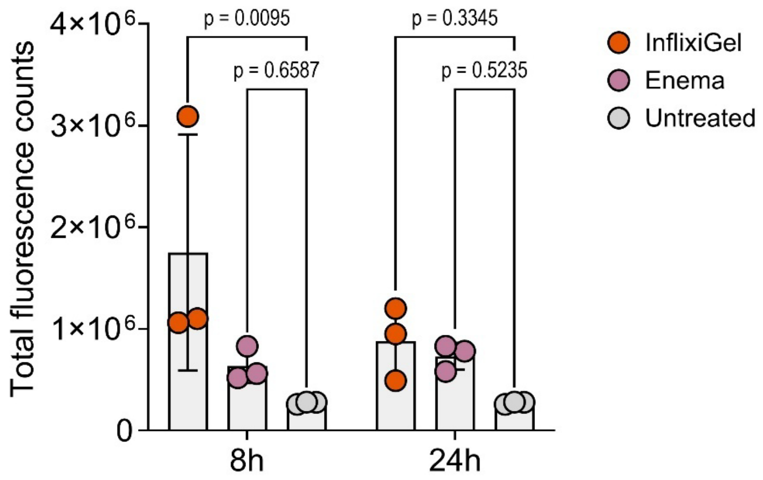


**Supplementary Figure 10.** Total fluorescence counts of untreated colon tissues or colon tissues at 8 and 24 h after administration of rectal formulations loaded with DiD dye (n=3).

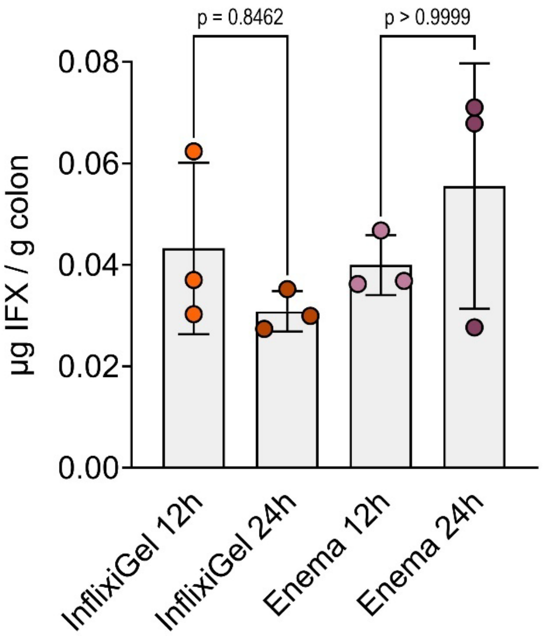


**Supplementary Figure 11.** Colonic concentration of IFX in healthy rats 12 and 24 h after rectal administration of InflixiGel or enema (n=3).

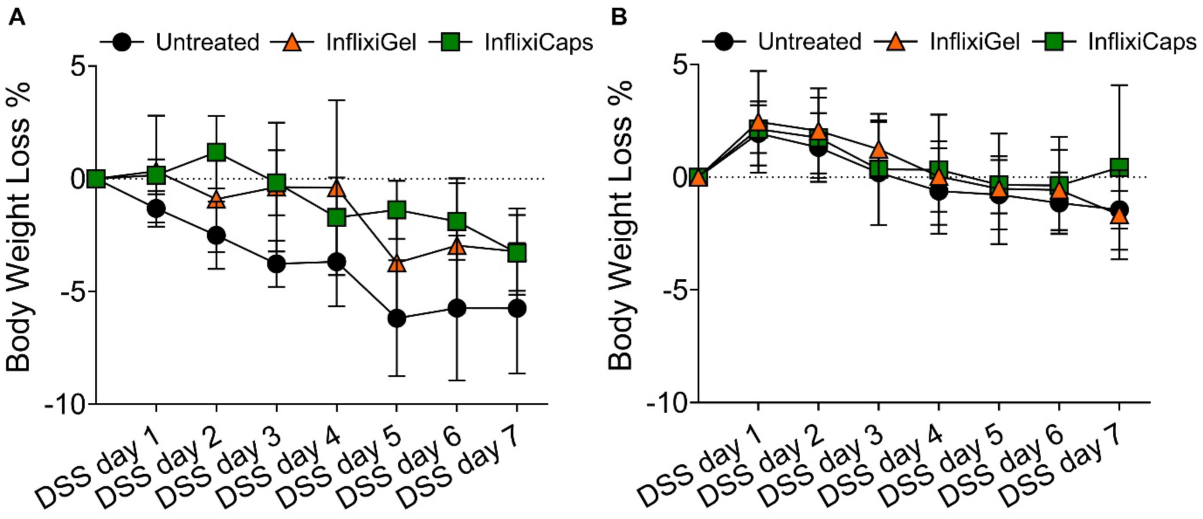


**Supplementary Figure 12.** Body weight loss of untreated rats and InflixiGel- and InflixiCaps-treated rats during seven days of (A) 4% DSS (w/v) cycle and (B) 6% DSS (w/v) cycle. Results are shown as mean ± SEM (n=6).

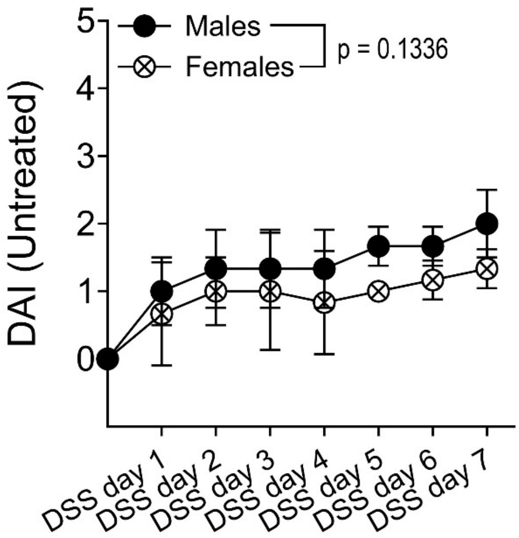


**Supplementary Figure 13.** Disease activity index (DAI) of untreated males and females during the seven days of 4% DSS (w/v) cycle. Results are shown as mean ± SD (n=3). Statistical significance was evaluated using Mixed-Effects model and the adjusted p-value corresponds to the comparison on the $7^{th}$ day.

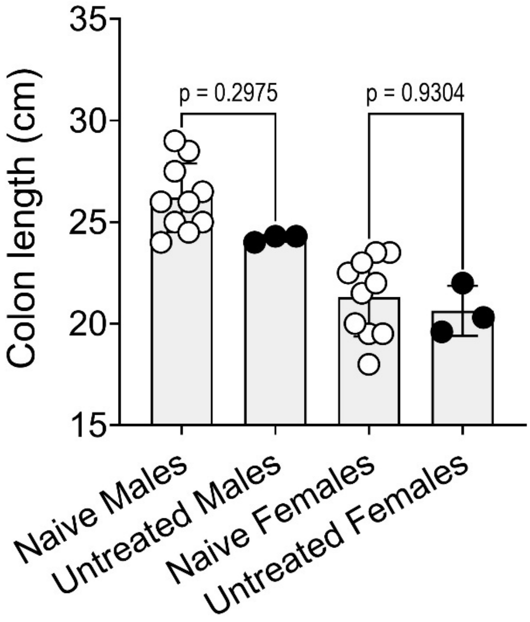


**Supplementary Figure 14.** Colon length of naïve and untreated males and females after the seven days of 4% DSS (w/v) cycle. Results are shown as mean ± SD. Statistical significance was evaluated using one-way ANOVA with Tukey's post hoc analysis. The adjusted p-values are reported above the comparison.

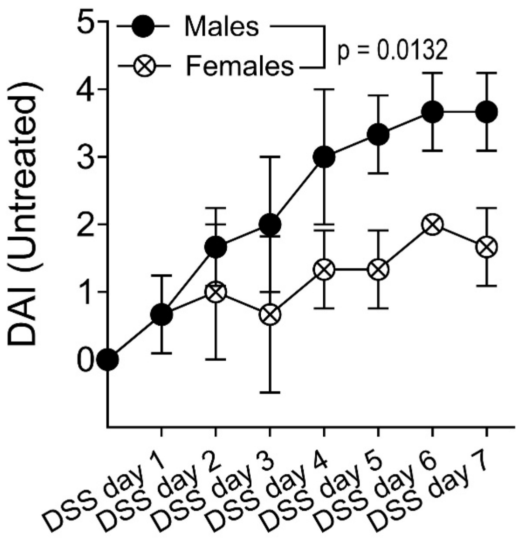


**Supplementary Figure 15.** Disease activity index (DAI) of untreated males and females during the seven days of 6% DSS (w/v) cycle. Results are shown as mean ± SD (n=3). Statistical significance was evaluated using Mixed-Effects model and the adjusted p-value corresponds to the comparison on the 7$^{th}$ day.

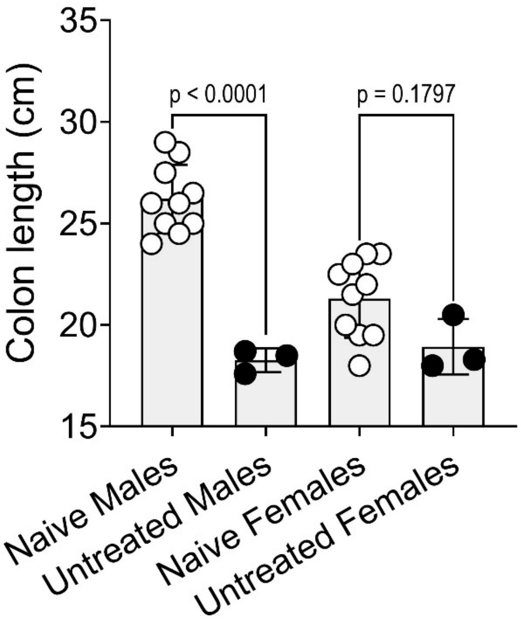


**Supplementary Figure 16.** Colon length of naïve and untreated males and females after the seven days of 6% DSS (w/v) cycle. Results are shown as mean ± SD. Statistical significance was evaluated using one-way ANOVA with Tukey’s post hoc analysis. The adjusted p-values are reported above the comparison.

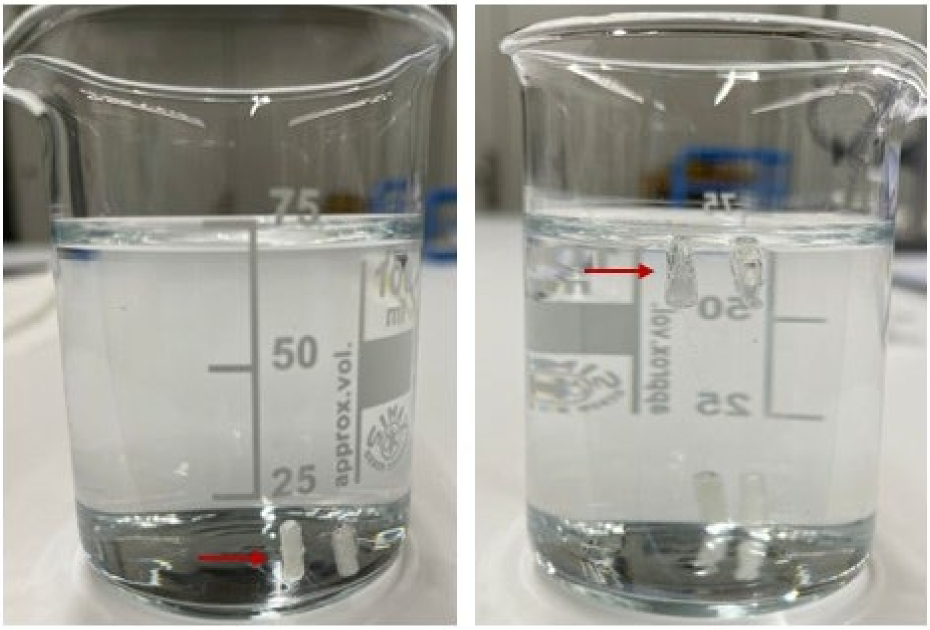


**Supplementary Figure 17.** Behavior of InflixiCaps (left) and PowderCaps (right) in solution, representative images. Capsules are indicated by arrows.

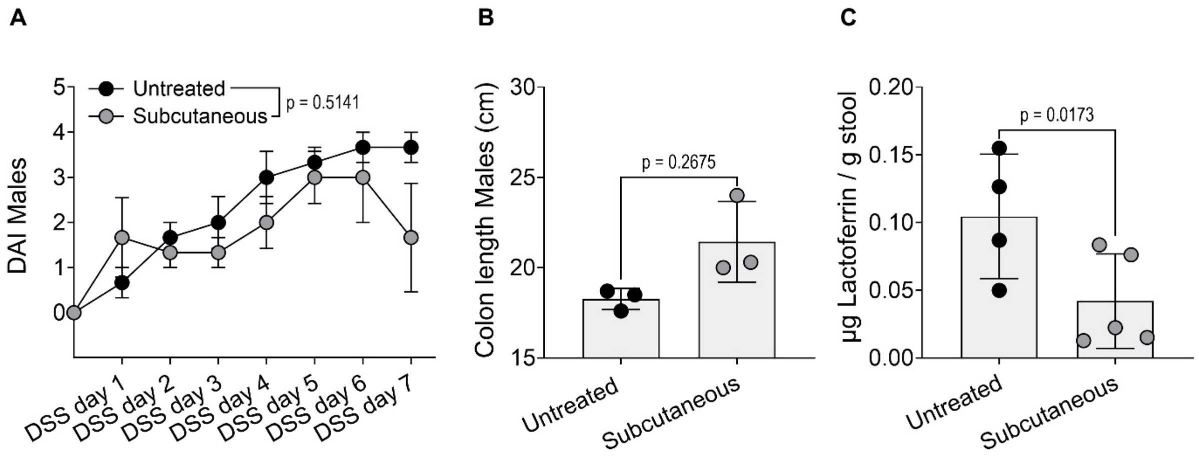


**Supplementary Figure 18.** Disease related endpoints of untreated rats and rats treated with subcutaneous injection of IFX after the seven days of 6% DSS (w/v) cycle. (A) Disease activity index (DAI), (B) colon length and (C) fecal lactoferrin concentration. Results are shown as mean ± SEM (n=3) in A. Statistical significance was evaluated using Linear Mixed-Effects. Results are shown as mean ± SD in B and C. Statistical significance was evaluated using nonparametric Mann-Whitney test and the adjusted p-values are reported above the comparison.

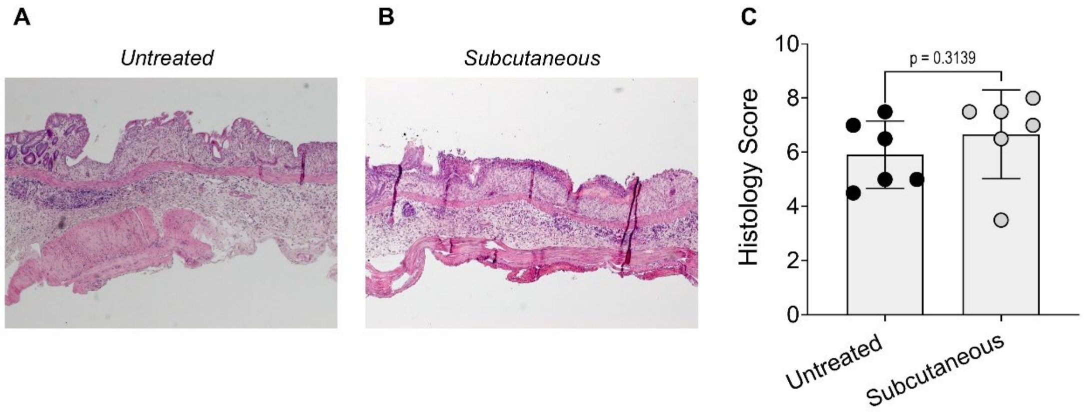


**Supplementary Figure 19.** Representative histology images of (A) untreated rats and (B) rats treated with subcutaneous injection of IFX after the seven days of 6% DSS (w/v) cycle, and (C) their histology score. Results are shown as mean ± SD. Statistical significance was evaluated using nonparametric Mann-Whitney test. The adjusted p-value is reported above the comparison.
